# Interpersonal alignment in infra-slow EEG rhythms and bodily signals anticipates mutual recognition

**DOI:** 10.1101/2025.05.29.656704

**Authors:** Nicolás Gravel, Simon Guendelman, Diego Vidaurre

**Affiliations:** Cognitive Neuroimaging Unit, Institut National de la Santé et de la Recherche Médicale, Commissariat à l’Energie Atomique et aux Énergies Alternatives, Centre National de la Recherche Scientifique, Université Paris-Saclay, NeuroSpin, Paris, France; Clinical Psychology of Social Interaction, Institute of Psychology, Humboldt-Universität zu Berlin, Germany; Berlin School of Mind and Brain, Humboldt-Universität zu Berlin, Germany; NeuroCure Cluster of Excellence, Charité - Universitätsmedizin Berlin, Germany; Centre de Recerca Matemàtica, Barcelona, 08193, Spain; Oxford Centre for Human Brain Activity, Wellcome Centre for Integrative Neuroimaging, Department of Psychiatry, University of Oxford, Oxford, OX3 7JX, United Kingdom; Department of Clinical Medicine, Center of Functionally Integrative Neuroscience, Aarhus University, 8000 Aarhus, Denmark

**Keywords:** Infra-slow EEG oscillations, EEG hyper-scanning, brain-body interactions, perceptual crossing, social coordination

## Abstract

Across multiple timescales, neural rhythms coordinate cognitive and physiological processes. Slower rhythms are increasingly recognised as the scale at which autonomic and interoceptive processes are folded into ongoing cognition, a brain–body view of allostatic regulation that is particularly relevant to social interaction. Examining these slow components of the electroen-cephalogram (EEG) during real-time interaction remains challenging. Here, we recorded infra-slow EEG alongside respiratory, cardiac and skin conductance signals during the perceptual crossing experiment, a social interaction and EEG hyperscanning paradigm. We assessed inter-personal alignment by contrasting trials in which dyads achieved mutual recognition with trials in which they did not, which separates genuine social alignment or recognition from a shared anticipatory context. Dyads achieving mutual recognition showed enhanced inter-participant alignment of ∼0.1 Hz infra-slow EEG power relative to those that did not, after accounting for the pre-movement motor potential. Alignment also differed between these trials in respiratory depth, the cardiac vagal index and skin conductance. These findings suggest that infra-slow EEG rhythms around ∼0.1 Hz may provide a temporal scaffold for the interpersonal coordination of brain and body, and could serve as markers for investigating the coupled processes underlying shared understanding and mutual recognition.

## 1 Introduction

Social interaction is not only a matter of representing another person internally, but also of coordinating perception, action, and bodily state with them in real time [1]. Across many approaches to social cognition, this has led to growing interest in social alignment: the idea that shared understanding emerges, at least in part, through the temporal organisation of coupled activity between interacting partners [2, 3]. Such alignment can be expressed behaviorally, physiologically, and neurally, and is believed to be especially relevant in situations where social meaning must be established online through ongoing mutual adjustment [3]. Rather than functioning as independent, parallel mechanisms for individual cognition, the coupling between neural and physiological systems may better be described as a system-wide network for cognitive and physiological regulation [4]. In social contexts, this regulation may extend across individuals as a form of interpersonal alignment or coregulation [5].

Social interaction and cognition related processes are supported by neuronal activity across multiple spatial and temporal scales [6, 3, 7], ranging from circuit-level field potentials to system-wide communication between brain and body [8]. Faster rhythms are often linked to moment-to-moment sensory processing, attention, and action, whereas slower fluctuations are increasingly understood as providing a broader temporal scaffold for cognition and bodily allostatic regulation [9, 5, 10, 11]. This slower end of the spectrum is especially relevant for socially situated behavior, because interaction unfolds not only in milliseconds but also over seconds-long windows in which expectation, arousal, anticipation, and mutual orientation are gradually organised [12, 1, 3].

Among these slow dynamics, infra-slow EEG fluctuations have attracted increasing interest as a possible bridge between brain activity and bodily regulation [13, 14, 15]. Activity below approximately 0.2 Hz has been associated with changes in arousal, interoceptive processing, and slow autonomic dynamics [10, 16, 11], and a rhythm around *∼*0.1 Hz is of particular interest because it lies near the timescale of prominent physiological fluctuations linked to respiration and heart–autonomic regulation, and ongoing allostatic control [17, 18, 19]. For example, meta-state transitions in the BOLD signal [20] tend to exhibit a periodicity of around 10 seconds and align with salient narrative shifts [12]. This suggests a relationship between slow body-wide physiological dynamics and meaningful synchronic changes in cognition and context [5, 16] —one that might be specially relevant for co-regulation of bodies across partners and that may be detectable by other means, including EEG [21]. From this perspective, infra-slow EEG is not simply a background feature of neural activity, but a candidate temporal scaffold through which brain and body are coordinated during engaged cognition [11].

This perspective is especially relevant for social interaction. If successful social interaction depends partly on each partner’s embodied allostatic regulation, this should be reflected, not only in overt behavior, but also in the alignment of slow neural and autonomic dynamics across partners [2, 22, 3]. In other words, mutual recognition may be supported by a shared brain-body state that unfolds over seconds and helps structure the timing of perception, action, and response among individuals [11, 12, 1]. Across social species, these processes and mechanisms are believed to be well preserved as constituents of basic forms of social interaction [23]. In humans, this suggests that slower physiological and neural components may be especially relevant when processes like social understanding or mutual recognition emerge gradually through shared context rather than through isolated, momentary stimulus processing [3]. This predicts that social alignment could be observed not only in behavior, but also in infra-slow EEG and in peripheral physiological signals closely tied to autonomic state, such as respiration and electrodermal activity [5, 24]. Hyperscanning, the simultaneous recording of brain activity from interacting partners, has established inter-brain synchronisation during social interaction as a marker for social alignment [25, 26]. This method has mostly examined faster oscillatory bands, so it remains unknown whether infra-slow activity also aligns between partners, and whether this alignment extends to the peripheral signals to which infra-slow activity is linked.

To examine this possibility, we studied dyads engaged in the perceptual crossing experiment (PCE), a minimalist social interaction paradigm in which participants attempt to detect one another in a shared virtual environment [27, 22, 28]. The PCE is designed to facilitate the systematic analysis of real-time social interaction by employing a minimalist approach [27, 22]. Within this computer-mediated interaction, participants report becoming aware of each other’s presence, the onset of which they mark by providing a button press. This allows to investigate whether slow brain-body dynamics change around socially meaningful events, and whether such dynamics differ between dyads who achieve mutual recognition and those who do not [28, 3].

We therefore investigated three main research questions. First, does infra-slow EEG, and in particular the *∼*0.1 Hz component, reflect meaningful contextual transitions across the structure of the task? Here we build on previous work linking a *∼* 0.1 Hz infra-slow EEG rhythm to autonomic and interoceptive fluctuations [29, 10, 13, 15, 24]. Second, does interpersonal alignment in infra-slow EEG reflect genuine mutual recognition rather than the common anticipatory structure imposed by a shared protocol? Because both partners share the same protocol structure and trial timing, we test whether such alignment is specifically enhanced in dyads that achieve mutual recognition relative to those that do not. Third, does within-participant coupling between infra-slow EEG and bodily signals reorganise around the moment of recognition? Taken together, these questions test whether interpersonal alignment can be tracked in slow brain-body dynamics and whether infra-slow EEG provides a useful window onto the embodied processes that support mutual recognition [5, 11, 3].

## 2 Methods

### 2.1 Participants

Data for this study were drawn from the ECSU–PCE dataset [28], collected at the Embodied Cognitive Science Unit (OIST, Japan) between August and December 2023. The full dataset comprises 32 pairs (64 participants, 39 identifying as female) aged 18–68 years (mean ± SD: 28.6 ± 8.9). Eligibility was determined by a preexperiment screening form that excluded psychiatric or neurological conditions, psychoactive medication, and allergic reactions to alcohol-based EEG preparation products. The study was approved by the OIST Human Subject Research Review Committee (HSR-2023-013), and all participants gave written informed consent. Our analyses used 31 of the 32 experimental sessions (one was excluded due to acquisition failure). The dataset is openly available (see Data and materials availability).

### 2.2 Experimental setup

We utilized the haptic perceptual crossing experiment (PCE) [27, 22, 28], a paradigm in which two participants, seated in separate rooms and with no visual or auditory contact, attempt to recognize each other’s presence in a one-dimensional circular virtual space mediated by haptic feedback. Each participant navigated the space by rotating a handle on a custom haptic interface; a brief vibration signalled any overlap with one of three possible objects: the partner’s avatar, a motionless static object, or a “shadow” object tracking the partner’s avatar at a constant offset (and therefore never corresponding to mutual recognition). Crucially, only partner–partner crossings produce a bidirectional vibration (both participants feel the event simultaneously), whereas crossings of the static and shadow objects are unidirectional. Participants were asked to press a button (making a click) whenever they believed they were interacting with the partner. Clicks were not perceivable by the other participant. Each trial thus yields one of three outcomes per participant: recognition leading to mutual recognition (joint success, *JS*, both participants click the other), one-sided recognition (single success, *SS*, only one participant clicks), or no recognition / wrong click (non-*JS* includes *SS* and *W* in the present analyses). The experimental protocol comprised (i) an initial 3-minute eyes-closed resting state, (ii) three training trials (two eyes-open, one eyes-closed) to familiarise participants with the device, and (iii) three batches of six 1-minute experimental trials (18 trials total) performed with eyes closed. Additional 3-minute resting-state recordings were acquired after trials 6, 12 and 18, yielding four resting states per session. After each trial, participants completed a short questionnaire on their subjective experience, including a Perceptual Awareness Scale (PAS) rating of the clarity with which they had experienced the partner just before the click. The full protocol, PCE device, recruitment procedure, and questionnaires (Big Five BFI-2; state-anxiety; subjective-experience items) are described in detail in the ECSU–PCE dataset paper [28].

### 2.3 Data acquisition and preprocessing

Electroencephalography (EEG) was acquired with a 64-channel active actiCAP snap set connected to a battery-operated BrainAmp DC amplifier (Brain Products GmbH, Gilching, Germany), with electrodes positioned according to the international 10–20 system. Recording and reporting followed published guidelines for EEG studies [30]. Signals were sampled at 1000 Hz. To preserve infra-slow EEG fluctuations, the amplifier was operated in DC-coupled mode with the acquisition cutoff set to 0.016 Hz–1000 Hz [13, 15, 31]. We note that this lowest-frequency setting still attenuates activity below *∼*0.08 Hz, which motivates restricting analysis to infra-slow EEG fluctuations around *∼*0.1 Hz (see Spectral parameterisation). Peripheral physiological signals were acquired synchronously on a battery-operated BrainAmp ExG amplifier (Brain Products) at 1000 Hz. Both amplifiers were recorded with BrainVision Recorder (v1.20.0502). They comprised the electro-cardiogram (ECG), respiration, recorded with a piezo-electric belt on the upper abdomen, and electrodermal activity (EDA, skin conductance). Electrode placements are given in the Supplementary Methods, and the full preparation procedure, device specifications, and data-collection timeline are documented in the ECSU–PCE dataset paper [28].

We then down-sampled the EEG data by a factor of 10 (from 1000 Hz to 100 Hz) and marked transient artefacts (“bad timestamps”) as 10 consecutive samples (1 s) for which the signal exceeded 5 × median absolute deviation (MAD) above or below the running median computed in a sliding 10 s window. Rejected samples were subsequently recovered using spherical spline interpolation [32], resulting in artefact-corrected signals. We did not apply independent component analysis to remove ocular artefacts. All task trials were performed with eyes closed, which limits blinks and saccades, and remaining transients were handled by this rejection step. To exploit the spatio-temporal structure of infra-slow EEG fluctuations at the protocol-level (across rest, pause and task), we spatially whitened the artefact-corrected signals obtained in the previous step using generalized eigenvalue decomposition (Supplementary Methods).

Each participant completed 18 trials, resulting in a total of 558 trials across all experiments. Of these, 206 trials had a button press from both participants between 15 and 45 s into the trial. A ±15 s epoch could therefore be extracted around each press (412 clicks: 302 *JS*, 41 *SS* and 69 *W* ). These 206 trials entered the EEG and respiratory analyses. The electrodermal signal was usable in 196 of them and the ECG in 169. For the protocol-level analysis (across rest, pause and task), we selected 62 participant-paired trials with a resting state period of at least 3 minutes, followed by a short interim period of 10 s before task onset, totalling within-participant 124 trials. Only trials that follow a resting-state recording contain the transition from rest to task, so this analysis was restricted to them.

### 2.4 Spectral parameterisation of infra-slow EEG fluctuations

To track transitions between task states at the protocol level, we decomposed the preprocessed EEG data into its spectral components using the multitaper method [33, 34, 35]. The multitaper method provides optimal estimates of spectral concentration by applying multiple orthogonal tapers (Slepian functions) to the signal, reducing variance while maintaining good frequency resolution [35]. For all the 124 protocol-level trials, the spectrum was computed across frequencies from 0.016 to 0.2 Hz in steps of 0.005 Hz using 3 Slepian tapers. A sliding time window of 60 seconds was applied with frequency smoothing of ±0.005 Hz with an overlap of 20 s. To ensure that the resulting spectrograms were comparable across participants, we normalized the spectral power by dividing each trial’s power by its temporal standard deviation, which allowed us to account for arbitrary differences across participants while preserving the relative dynamics of spectral power.

To separate the periodic component from the 1/f aperiodic background, we adapted the *fitting oscillations and one over f* (FOOOF) method [36], fitting the aperiodic background and two Gaussian peaks at 0.05 Hz and 0.1 Hz to the multitaper PSD. FOOOF models the PSD as the sum of a broadband aperiodic component (1/f background) and periodic oscillatory peaks, enabling quantitative comparison of spectral features across conditions. The two centre frequencies were pre-specified from earlier reports of infra-slow EEG fluctuations. Only the amplitudes and widths of the Gaussians were estimated, so the model does not select these frequencies from the data. Given the small number of frequency bins, this minimises overfitting and ensures a biologically meaningful parameterisation of the infra-slow EEG fluctuations of interest. The 1/f slope was modelled in log–log space and subtracted from the PSD at each time bin, and the residual PSD was fitted with the two Gaussians centred at 0.05 Hz and 0.1 Hz. Whereas FOOOF relies on Welch’s spectral decomposition method, we chose a small number of Slepian (discrete prolate spheroidal sequences, DPSS) tapers. This choice was motivated by the limited number of data points and cycles available in the infra-slow EEG range, where Welch’s segmentation and averaging can produce unstable and biased estimates. Slepian tapers provide optimal spectral concentration and reduced leakage, yielding more reliable power estimates for ultra-low frequencies with sparse data.

Because the 0.016 Hz hardware high-pass attenuates activity below *∼* 0.08 Hz [31], the 0.05 Hz Gaussian is retained as a descriptive spectral feature only; all maintext task-level inferential analyses (APIM and phase-at-click) are restricted to the 0.1 Hz component (0.08–0.12 Hz band). To assess the stability of the fits, the model was fitted separately for each channel and time point in five folds, each leaving out 20% of the recordings. We report the mean and standard deviation of the goodness of fit (*R*^2^) across channels, time points and folds.

### 2.5 Modelling interpersonal alignment in task-locked *∼*0.1 Hz EEG power

Informed by our initial exploratory spectral decomposition, we applied a zero-phase band-pass Butterworth filter (4th order, cutoff frequencies 0.08 and 0.12 Hz) to the artefact-corrected 100 Hz data. We then Hilbert-transformed the filtered EEG data across each 60-second trial. From the resulting analytical signal, we extracted the instantaneous phase and computed the instantaneous power as the squared absolute magnitude of the amplitude envelope for each of the 64 channels. Instantaneous power was further reduced to a single time series by extracting the first principal component after subtracting the temporal mean across channels. We used a single component because the model requires one outcome per participant and window, and because power in this band is largely shared across the scalp: the first component explained a median of 92% of the variance (interquartile range 84–98%), and its mean loadings had the same sign on all 64 channels. It therefore indexes a global fluctuation of infra-slow power. To avoid edge effects from filtering and the Hilbert transform, we selected trials with task responses (button clicks) occurring between 15 and 45 seconds into the trial, extracting epochs centered ±15 seconds around button clicks.

To quantify interpersonal alignment in the *∼*0.1 Hz infra-slow EEG band at the group level (*JS* vs *non-JS* ), we fitted a symmetric Actor–Partner Interdependence Model (APIM) [37, 38] implemented as a linear mixed-effects regression with restricted maximum likelihood (REML; Supplementary Methods). Its partner × condition term estimates, at each window, how mutual recognition modulates interpersonal alignment. The model was fitted independently at five contiguous 6-second windows centred at −12, −6, 0, +6 and +12 s relative to each button press. Per-window REML coefficients (*β*) and their asymptotic standard errors were used to compute one-sided (*JS* > *non-JS* ), uncorrected Wald p-values for the partner × condition interaction; statistical significance is reported directly from the LME, and in the figures the asterisks that also survive the Benjamini–Hochberg correction across the five windows of a panel (false discovery rate 0.05) are red colored. The APIM accounts for the dyadic non-independence of paired participants and for unequal trial counts across experiments via partial pooling. We call the windows before the button press anticipatory, a term that refers to their timing relative to the press (see Methodological considerations). To test whether each condition’s slope was press-locked beyond autocorrelation, we also evaluated it against an onset-shift permutation null (N=200). Statistics were combined across the five windows with Stouffer’s method.

Because the analysis is time-locked to button presses, the pre-movement motor potential — a slow cortical potential preceding voluntary movement [39] — constitutes a confound that can leak into the infra-slow band and inflate apparent alignment. To remove this confound, we regressed out a motor-potential template from all 64 channels before bandpass filtering. For each participant and trial, the press-aligned event-related potential (ERP) was computed over motor channels (C3, Cz, C4), baseline-corrected to [−15, −12] s, multiplied by a Tukey window, and the template, scaled by an ordinary least-squares (OLS) fit with an intercept, was subtracted from all 64 channels within ±15 s of the press, before bandpass filtering. This regression step is applied to EEG only; respiratory and electrodermal signals are not contaminated by cortical motor slow potentials and are therefore left unchanged. Every analysed trial contained a button press from both participants, in *JS* and *non-JS* alike, and each epoch was aligned to that participant’s own press, so the presence of a press does not distinguish the two outcomes.

We also measured how closely the two partners pressed in time. For each analysed trial, the response delay was the absolute difference between the two participants’ press times (follower minus leader), taken from the click time stamps and expressed in seconds. Trials were grouped by outcome, using the 206 trials of the 31 analysed sessions (*JS*, 151 trials; *non-JS*, 55 trials). Because delays are right-skewed, we summarise them by their median and interquartile range, together with the mean and standard deviation, and compared the two outcomes with a two-sided Mann–Whitney *U* test, which treats trials as independent.

### 2.6 Modelling interpersonal alignment in respiratory, cardiac and electrodermal signals

The respiratory belt signal was recorded with inverted polarity, so we flipped its sign before the inter-participant alignment, phase-at-click and phase-coupling analyses. The respiratory cycle was derived from the respiratory belt signal as a full-cycle index in [0, 1], where 0 marks the expiratory trough, 0.5 the inspiratory peak, and 1 the next trough (Supplementary Methods). To test whether responses favoured a phase of the respiratory cycle, we read the full-cycle index at each click and applied Rayleigh tests to the phase values, separately for each condition. The cleaning of the respiratory and ECG signals, the detection of respiratory peaks and troughs, the phase-completion estimator, the detection of R peaks and the ECG delineation used NeuroKit2 [40]. EDA was decomposed into tonic (skin conductance level) and phasic (skin conductance response) components. The same APIM model used for the EEG was then fitted to each physiological channel. To quantify the depth of breathing, we fitted the model to respiratory depth: the upper envelope of the squared running root-mean-square of the sign-corrected and cleaned belt signal, computed over 3 s, about one breath (Supplementary Methods). The interaction was tested one-sided (*JS* > *non-JS* ) for respiratory depth and in each direction for the electrodermal signals, uncorrected.

To include a measure of beat-to-beat cardiac regulation, the same model was fitted to the cardiac vagal index (defined below), computed over 5 s around each window centre, with the interaction tested one-sided (*JS* > *non-JS* ).

### 2.7 Within-participant brain–body coupling analysis

To assess the temporal correspondence between the infra-slow EEG fluctuations (*∼* 0.1 Hz) and physiological signals at the individual level, we examined brain–body coupling by pooling data from both participants within each trial. We fitted a within-participant linear mixed-effects model (Supplementary Methods) at the same five 6-second windows used for the APIM analysis, with REML estimation and Wald p-values for the physio × condition interaction, tested one-sided in each direction and uncorrected. Respiratory and electrodermal signals were processed identically to the dyadic analysis. To disentangle motor-cortex slow potentials from genuine brain–body coupling, the model was fitted both with raw EEG and with EEG from which the pre-movement motor potential template had been regressed out; in this analysis, the template was averaged across each participant’s trials. To include a measure of beat-to-beat cardiac regulation, we computed the cardiac vagal index [41, 40] from the R–R intervals of the ECG. This non-linear measure takes the geometry of the Poincaré plot, a scatter plot of each interval between heartbeats against the next, and turns it into a proxy for parasympathetic (vagal) control of the heart. Because the index needs several heartbeats, we computed it over 5 s around each window centre (Supplementary Methods).

### 2.8 Phase coupling between the respiratory and cardiac cycles

To test whether the coupling between respiratory and cardiac phase at the moment of the response differed with task outcome, we modelled the two phases jointly with a bivariate von Mises sine model [42]. For each button press, respiratory phase *ϕ*_*r*_ was 2*π* times the full-cycle respiratory index at the click. Cardiac phase at the click was built from the R peaks and T offsets of the ECG (Supplementary Methods) in four contiguous segments, each linear in time: systole, from the R peak to the T offset, occupied one quarter of the cycle, and diastole, from the T offset to the next R peak, the other three quarters, divided into early, mid and late thirds. The coupling parameter of the model, *λ*, is zero when the phases are independent (Supplementary Methods), and we took *Δλ* = *λ*_JS_ *−λ*_non-JS_ as the test statistic.

To obtain a null distribution that preserves the auto-correlation of the signals, we drew 200 sets of surrogate click times, independently for each click and uniformly between 15 and 45 s into the trial. We kept the observed condition labels, re-extracted both phases at the surrogate samples with the same procedure, and refitted the model. The two-sided *p* value of *Δλ* was the proportion of surrogates with |*Δλ*| at least as large as observed. To locate where in the phase plane the clicks gathered, we counted them in the 16 combinations of two quarter-cycle bins and tested each cell against 50 of the surrogate sets, with a correction across the cells (4 × 4 test; Supplementary Methods).

We fitted the same model, with the same surrogate procedure, to the phase of the infra-slow EEG paired with the respiratory and with the cardiac phase (Supplementary Methods; Supplementary Results).

## 3 Results

Our analyses addressed three questions framed around the contextual structure of the PCE. At the protocol level, each trial moves through three states — rest →eyes-open pause → task — that define a slowly varying behavioural and perceptual context. In the original protocol, the trials eligible for this level are those that directly follow a resting-state recording: the first trial of the second and third batches, that is, trials 7 and 13 of each session (rests were recorded after trials 6, 12 and 18). The first batch follows the training trials, and the last rest marks the end of the experimental session. These trials give the 62 participant-paired trials (124 participant trials) of the protocol-level analysis. At the task level, the critical event is the participant’s button press marking the subjective moment of recognising the other. First, we asked whether *∼* 0.1 Hz infra-slow EEG activity could be reliably characterised and whether it tracked protocol-level contextual transitions. Second, whether inter-participant alignment in the infra-slow EEG and in accompanying autonomic signals (respiratory depth, electrodermal activity) was sensitive to the task-level transition around the button press and to task outcome (mutual recognition versus not). Third, whether within-participant brain–body coupling around the response reorganised with task outcome.

### 3.1 Infra-slow EEG activity tracks contextual transitions at the protocol level

Figure 1 (left) presents our spectral-decomposition strategy. The multitaper PSD (Slepian/DPSS tapers) of infra-slow EEG activity in the 0.016–0.2 Hz range is dominated by a 1/f aperiodic background that conventional Fourier approaches cannot cleanly separate from narrow-band oscillatory content. To isolate the periodic component, we parameterised the PSD using a custom implementation of the *fitting oscillations and one over f* (FOOOF) method [36]. The parameterisation revealed a clear periodic peak at *∼* 0.1 Hz superimposed on the 1/f aperiodic background. Across channels, time points and folds, the 1/f fit had a mean *R*^2^ of 0.676 (SD 0.072) and the two-Gaussian fit a mean *R*^2^ of 0.549 (SD 0.067). Figure 1 is descriptive. In a separate test on the task epochs, power at 0.08–0.12 Hz exceeded the 1/f background (Supplementary Results).

**Fig. 1.**
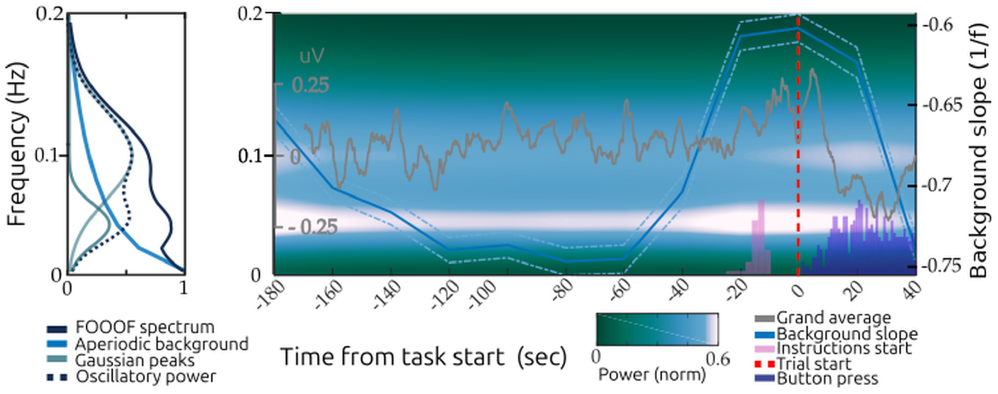
Spectral signature of the *∼* 0.1 Hz infra-slow EEG rhythm. Left: custom FOOOF parameterisation of the multi-taper PSD. Right: time-frequency representation of the infra-slow EEG, with the canonical task structure (rest → eyesopen pause → task) overlaid. Power is averaged over the 64 channels and the 124 protocol-level recordings. Time 0 marks the end of the 3-minute resting period.

Figure 1 (right) summarises the FOOOF results at the protocol level. The 1/f slope steepens during eyes-closed rest, flattens during the eyes-open pause, and flattens further during the task, where a distinct narrow-band peak at *∼*0.1 Hz emerges. The 0.08–0.12 Hz band emerged during the task and was suppressed during rest and pause. The per-state histograms in Fig 1 aggregate this pattern across trials. Together, these results establish a *∼*0.1 Hz narrow-band component of the infra-slow EEG that is sensitive to protocol-level contextual transitions.

### 3.2 Inter-participant alignment anticipates mutual recognition

Having isolated the *∼* 0.1 Hz infra-slow EEG component, we asked whether this component and accompanying autonomic signals aligned between paired participants around the button press, which is the key task-level contextual transition. Pairs were grouped by outcome into Joint Success (*JS*, both partners recognised each other) and non-Joint Success (*non-JS*, pooling single-success and wrong responses).

The analyses of *∼* 0.1 Hz infra-slow EEG power presented below was derived from the first principal component of the 64-channel power, in order to summarise global, mean-field tendencies across the whole brain (Methods). Partners pressed closer together in time when they recognised each other: the delay between the two presses of a trial had a median of 4.9 s (interquartile range 1.2–9.1 s; mean 6.6 s, SD 6.5 s) in *JS* trials (*n* = 151) and 8.5 s (3.6–14.4 s; mean 10.0 s, SD 6.9 s) in *non-JS* trials (*n* = 55). The difference was significant (two-sided Mann–Whitney *U* = 2782, *p* = 0.0003; rank-biserial correlation 0.33).

#### 3.2.1 Inter-participant alignment in the 0.1 Hz infra-slow EEG

Inter-participant alignment in the instantaneous power of the *∼* 0.1 Hz component around button presses was estimated with a symmetric Actor–Partner Interdependence Model (APIM) fitted by restricted maximum likelihood (Fig 2A). The outcome was the first principal component of the 64-channel power (see Methods). At each of five 6-second windows, the model included partner, condition and their interaction as fixed effects, with a by-dyad random intercept and partner slope. Prior to fitting, we regressed out the pre-movement motor potential from the EEG to remove the slow motor confound time-locked to the button press. The partner × condition APIM slope was significantly larger in *JS* than *non-JS* pairs in the windows surrounding mutual recognition (Stouffer Z = 2.00, *p* = 0.023) (Fig 2A). At *t* = − 6 s (*JS β* = 0.17, 95% CI 0.02 to 0.32; *non-JS β* = − 0.13, 95% CI − 0.34 to 0.07; difference 0.31, 95% CI 0.09 to 0.53) the difference also survived the false discovery rate correction across the five windows (*q* = 0.016) (Fig 2A). Estimates and confidence intervals for every window of Figures 2 and 3 are given in Table S1. The effect held when the first component was computed without frontopolar, anterior-frontal and rim channels (Supplementary Results). *JS* slopes were press-locked beyond autocorrelation chance (onset-shift null, Z = 3.14, *p* < 0.001), most clearly in the windows at *t* = − 6 and 0 s (2.6 and 1.9 SDs above the null mean). *Non-JS* slopes were not distinguishable from the null (Z = 0.30, *p* = 0.38). Figure S1 reports a dyad-level ordinary least-squares fit used to assess trial-count balance across task outcomes and dyads, ruling out data imbalance as a driver of the task-level effects.

**Fig. 2.**
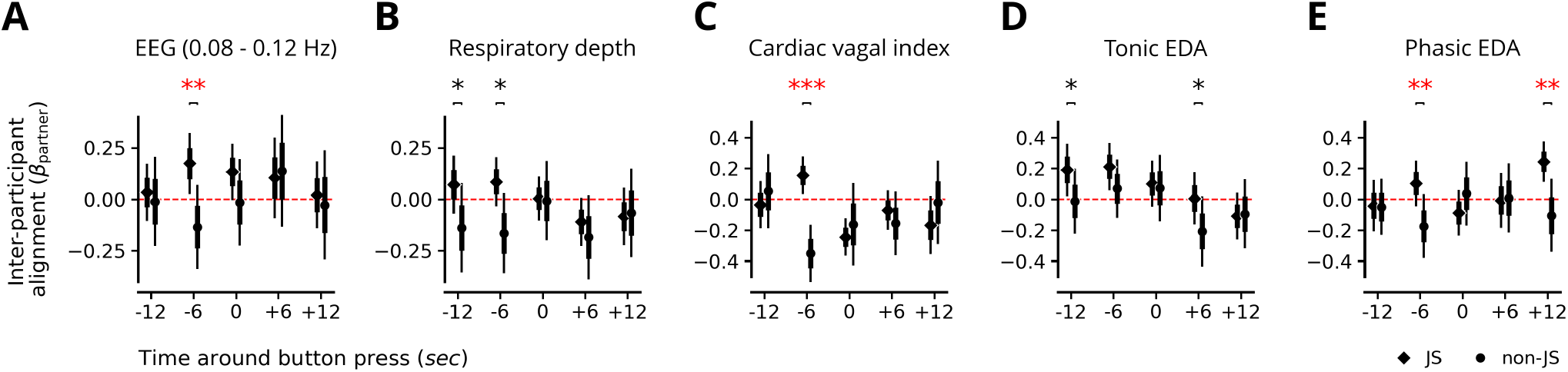
Increased alignment in infra-slow EEG power and autonomic signals around task response anticipates mutual recognition. Alignment at five windows centred on button presses, for Joint Success (*JS*) and non-Joint Success (*non-JS*) pairs. Alignment is the partner slope (*β*_partner_) of the Actor–Partner Interdependence Model (APIM; Supplementary Methods). Diamonds (*JS*) and circles (*non-JS*), estimates at each window; thin bars, 95% confidence intervals; thick bars, standard errors; red dashed line, zero. Asterisks: one-sided Wald test of *JS* > *non-JS*; * *p* < 0.05, ** *p* < 0.01, *** *p* < 0.001; red asterisks also survive the Benjamini–Hochberg correction across the five windows of the panel (false discovery rate 0.05). In D and E the interaction is tested in each direction, and the correction uses twice the smaller one-sided *p*. **(A)** Instantaneous spectral power of the 0.08–0.12 Hz band, after regressing out the pre-movement motor potential (N=151 *JS* and N=55 *non-JS* trials). **(B)** Respiratory depth (N=151 *JS* and N=55 *non-JS* trials). **(C)** Cardiac vagal index, computed over 5 s around each window centre (N=121 *JS* and N=48 *non-JS* trials). **(D)** Tonic skin conductance level. **(E)** Phasic skin conductance response, as one 5 s mean per window (N=145 *JS* and N=51 *non-JS* trials for D and E).

**Fig. 3.**
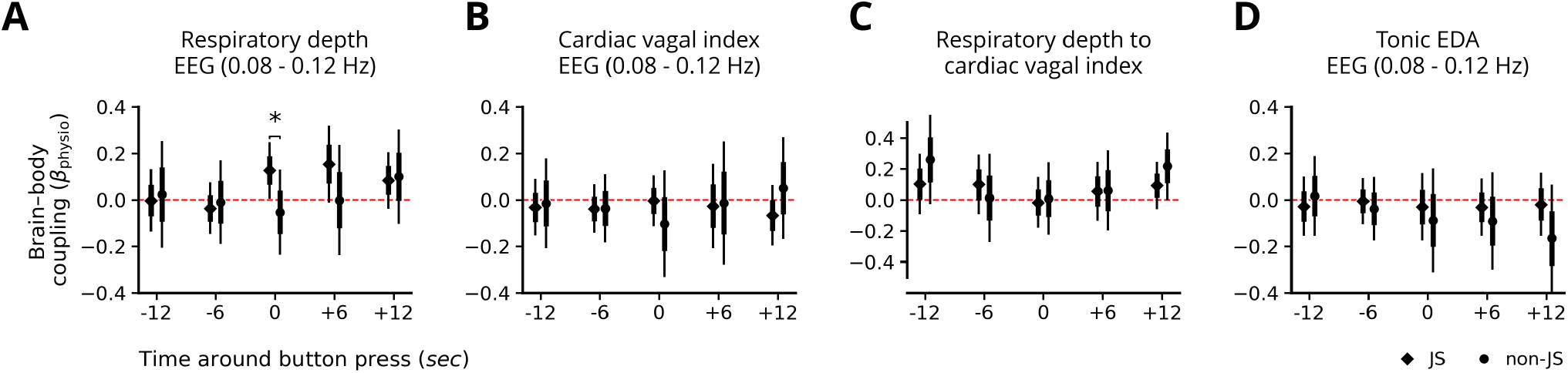
Similar infra-slow dynamics in brain-body coupling during successful and unsuccessful recognition. Brain– body coupling from the within-participant linear mixed-effects model at five windows centred on button presses, for Joint Success (*JS*, diamonds) and non-Joint Success (*non-JS*, circles) participants. Coupling is the slope of the predictor named first in each title (*β*_physio_; Supplementary Methods). Markers, estimates; thin bars, 95% confidence intervals; thick bars, standard errors; red dashed line, zero. Asterisk: one-sided test of the interaction in each direction; * *p* < 0.05; a red asterisk would also survive the Benjamini–Hochberg correction across the five windows of the panel (false discovery rate 0.05, on twice the smaller one-sided *p*); none does. **(A)** Respiratory depth as the predictor of the motor-regressed EEG (N=302 *JS* and N=110 *non-JS* clicks). **(B)** Cardiac vagal index as the predictor of the motor-regressed EEG, computed over 5 s around each window centre (N=242 *JS* and N=96 *non-JS* clicks). **(C)** Respiratory depth as the predictor of the cardiac vagal index, with no EEG (the clicks of B). **(D)** Tonic skin conductance as the predictor of the motor-regressed EEG (N=290 *JS* and N=102 *non-JS* clicks).

#### 3.2.2 Inter-participant alignment in respiratory depth, the cardiac vagal index and skin conductance

We next extended the dyadic APIM to autonomic signals recorded concurrently during the task: respiratory depth (Fig 2B) and the cardiac vagal index (Fig 2C). Alignment was evaluated at five time windows centred on the button press (−12, −6, 0, +6, +12 s).

Alignment of respiratory depth was stronger in *JS* than *non-JS* before the response, at *t* = − 12 s (*p* = 0.035) and *t* = − 6 s (*p* = 0.016; *q* = 0.088 and 0.078 after the false discovery rate correction across the five windows), and did not differ at the response itself (Fig 2B). Alignment of the cardiac vagal index was stronger in *JS* than *non-JS* at *t* = − 6 s (*JS β* = 0.16, 95% CI 0.03 to 0.28; *non-JS β* = − 0.35, 95% CI − 0.53 to −0.17; one-sided *p* < 0.001, *q* < 0.001 after the false discovery rate correction across the five windows), and did not differ at the other windows (smallest one-sided *p* = 0.25; Fig 2C).

To evaluate whether dyadic alignment extended to sympathetic arousal, we also measured electrodermal activity. Partner alignment in tonic and phasic skin conductance showed outcome-dependent temporal coordination (Fig 2D, E), confirming that social co-regulation engages peripheral sympathetic as well as parasympathetic dynamics.

Together, these results indicate that mutual recognition is accompanied by coordinated inter-participant alignment of autonomic dynamics, complementing the dyadic alignment of the *∼*0.1 Hz infra-slow EEG.

### 3.3 Within-participant brain–body coupling did not differ reliably with task performance

To address the third question, we fitted a within-participant linear mixed-effects model at each of the five time windows around the button press, for each EEG variant (raw versus motor-regressed; Supplementary Methods). The physiological signal × condition interaction indexes performance-dependent brain–body coupling (Fig 3).

Figure 3A shows the same model for respiratory depth and motor-regressed EEG. Coupling was larger in *JS* than *non-JS* at *t* = 0 (one-sided *p* = 0.047, uncorrected), a difference that did not survive the false discovery rate correction across the five windows (*q* = 0.48), and did not differ at the other windows (smallest one-sided *p* = 0.123).

For the cardiac vagal index, a measure of heart rate variability from the Poincaré plot of successive beats, coupling with motor-regressed EEG did not differ be-

tween *JS* and *non-JS* at any window (Figure 3B; smallest one-sided *p* = 0.180). Turning the model around, with respiratory depth as the predictor of the cardiac vagal index and no EEG, coupling did not differ at any window either (Figure 3C; smallest one-sided *p* = 0.17). Coupling with tonic EDA did not differ at any window either (Figure 3D; smallest one-sided *p* = 0.130; Supplementary Results).

### 3.4 Phase of the *∼* 0.1 Hz wave and of the respiratory cycle at the response

Beyond amplitude alignment, we asked whether task responses occurred at a preferred phase of the *∼* 0.1 Hz wave (Fig S2A, middle). Pooled across all 64 channels, phase distributions deviated from uniformity in both conditions and concentrated towards the peak of the wave in *JS* and after its trough in *non-JS* (*JS* : *z* = 8.8, *p* < 0.001, mean phase 161°, mean resultant length 0.02; *non-JS* : *z* = 14.5, *p* < 0.001, mean phase 82°, mean resultant length 0.05; both conditions peaked in the 90–180° bin). Because channels at one click are not independent, this result describes a global, scalp-wide tendency and not the phase of individual responses. With one value per click, phases did not deviate from uniformity in either condition (Supplementary Results), so we do not interpret the pooled tendency as a preferred phase of individual responses. Angles run from the wave trough (0°) through its peak (180°) to the next trough (360°). These phase values should be interpreted in light of the systematic *∼* 10° positive bias of the Hilbert phase-extraction approach documented in the phase-accuracy benchmark (see Supplementary Methods).

We next examined the within-participant timing of responses within the respiratory cycle (Fig S2B). Rayleigh tests applied separately to each condition showed that *JS* responses concentrated in the second half of expiration, while *non-JS* responses concentrated in the first half of expiration; both distributions were individually non-uniform (*JS* : *z* = 10.3, *p* < 0.001, mean phase 294°, mean resultant length 0.18; *non-JS* : *z* = 4.4, *p* = 0.012, mean phase 258°, mean resultant length 0.20). Phase runs from the expiratory trough (0°) through the inspiratory peak (180°) to the next trough (360°).

We next asked whether the respiratory and cardiac cycles were coupled at the click (Figure 4). Respiratory– cardiac coupling did not differ from the null in either condition (*λ* = +0.131, *p* = 0.378 for *JS* ; *λ* = +0.299, *p* = 0.353 for *non-JS* ) or between conditions (*Δλ* = − 0.169, *p* = 0.647), and no cell of the 4 × 4 test departed from chance (smallest uncorrected *p* = 0.059).

**Fig. 4.**
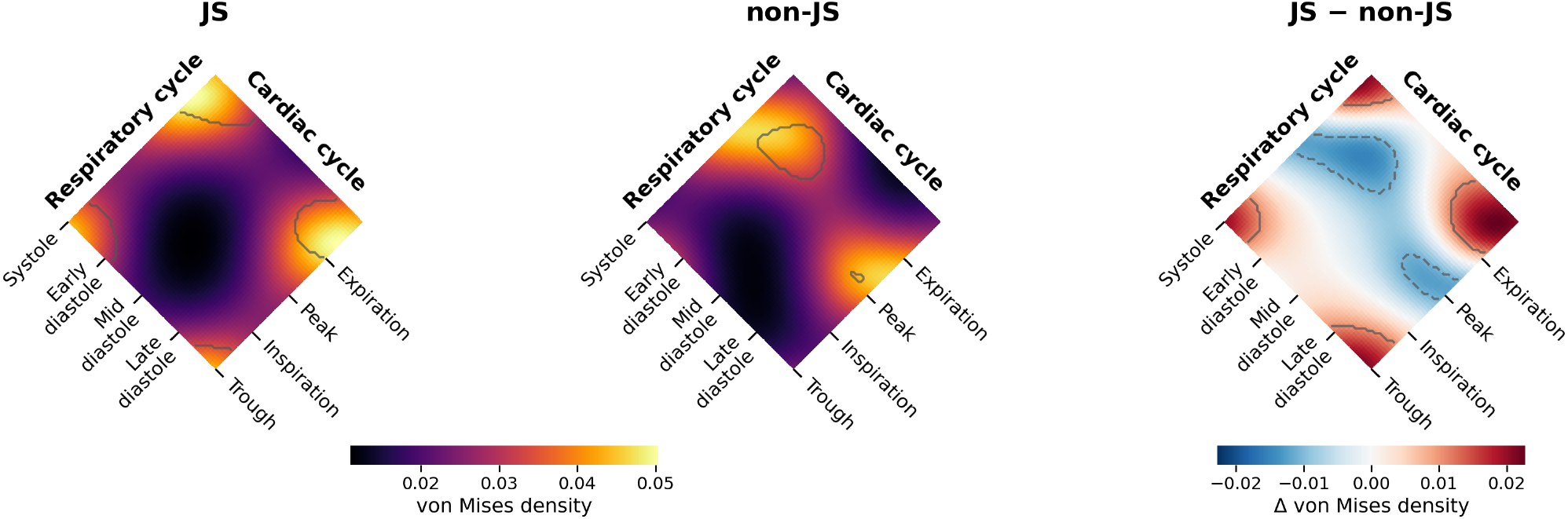
Phase coupling between the respiratory and cardiac cycles at the click. Fitted bivariate von Mises densities of respiratory cycle phase and cardiac phase for *JS* (N=270 clicks), *non-JS* (N=103 clicks) and their difference, as labelled above each column. The two densities share one colour scale. Curved outlines mark clusters above the 85th percentile of the onset-shift null: in the *JS* and *non-JS* columns, clusters of density above the mean of the null; in the difference column, clusters of the difference (solid, positive; dashed, negative). Black outlines, *p* < 0.05 in the cluster test; grey outlines, not significant (all). No cell of the 4 × 4 test, in which clicks were counted in the 16 combinations of two quarter-cycle bins and compared with the onset-shift null, held more clicks than chance or a larger share of *JS* than of *non-JS* clicks, so no square is outlined. Labels and tick marks give the position of each landmark on its axis: the trough and peak of the respiratory cycle with mid-inspiration and mid-expiration between them, and the centre of each cardiac segment. Cardiac phase runs through systole and early, mid and late diastole (Methods); because systole fills a larger share of each beat than of the axis, density gathers in systole even when clicks are spread evenly in time.

Along the respiratory axis, the density peaked in late expiration for *JS* and early expiration for *non-JS* clicks, as in Fig S2B. Along the cardiac axis, it was highest in systole in both conditions. Systole fills about 40% of each beat but a quarter of the axis, so this reflects the construction of the phase and not a preference for systole: against the chance expected from the time spent in each segment, no segment held more clicks than expected in either condition (Fig S2C; smallest one-sided *p* = 0.13). The same model applied to the phase of the infra-slow EEG, paired with the respiratory and with the cardiac phase, is reported in the Supplementary Results (Fig S3): coupling with the respiratory phase did not differ significantly with task outcome, and coupling with the cardiac phase differed between conditions (*Δλ* = +0.669, *p* = 0.040, uncorrected).

Our results demonstrate inter-participant alignment in *∼* 0.1 Hz infra-slow EEG power during social coordination. This alignment extends to autonomic physiology: respiratory depth, the cardiac vagal index and both components of the electrodermal signal align between participants. Within individuals, coupling between the infra-slow EEG and autonomic signals did not differ reliably with task outcome. Overall, our results point to interpersonal alignment of brain and body at infra-slow timescales around mutual recognition.

## 4 Discussion

Neuronal rhythms operate across multiple temporal scales, forming a complex basis for cognitive and physiological processes underlying social interaction behaviours. While fast oscillations have been extensively studied for their role in perception and attention [43], slower rhythms may provide a fundamental scaffold for integrating the brain’s various network dynamics [44, 6, 29] with allostatic bodily processes [5, 10] over the longer timescales necessary for social interaction processes [1, 3]. In our data, infra-slow EEG fluctuations and autonomic signals aligned between partners around mutual recognition. In what follows we discuss the major findings of our study, their implications for understanding the role of infra-slow EEG activity in social cognition, and the methodological considerations involved.

### 4.1 The role of infra-slow rhythms during social coordination

Our protocol-level analysis revealed a sequence of transitions in infra-slow EEG fluctuations that tracked meaningful changes in behavioural state, from rest to task. The sequential transition from aperiodic-dominated rest, through a flattened pause period, to a clear *∼* 0.1 Hz task-related peak, revealed by our protocol-level analyses (Fig 1), is consistent with the view that infra-slow rhythms provide a temporal scaffold for integrating brain and bodily dynamics during social interaction, and could reflect a number of mechanisms, such as the combined effect of slow readiness potentials during task engagement [45, 46], the accumulation of perceptual evidence [47, 48], adding to emerging evidence that infra-slow fluctuations correspond to different network configurations and cognitive states [49, 50, 51].

At the task-level, we linked the results obtained from our spectral parameterisation and dyadic APIM to task response outcomes. To examine inter-participant alignment of brain rhythms, we analysed the temporal relationship between infra-slow EEG fluctuations in participant pairs using absolute Hilbert envelopes as a proxy for spectral power. When comparing spectral power in relation to task responses (time-locked to button presses), our analysis revealed significantly stronger interpersonal alignment in the 0.08–0.12 Hz band, for participants who successfully recognised each other compared to those who did not (Fig 2A). Alignment was strongest in the windows surrounding mutual recognition (approximately 3–9 s before button press), as established by the per-window APIM linear mixed-effects model after regressing out the pre-movement motor potential (see Methods). Because this motor potential is common to every button press, removing it ensures that the residual interpersonal alignment reflects genuine social coordination rather than the shared temporal structure of the task (e.g. button press). The two partners’ presses within a trial were a median of 4.9 s apart in *JS* and 8.5 s apart in *non-JS* trials, and each epoch was aligned to its own participant’s press, so the windows we compared were typically offset in trial time by several seconds. The alignment we report therefore describes a shared, press-locked pattern of infra-slow power that differed with outcome, and not synchrony in the sense of co-fluctuation at the same moment; the onset-shift null, which showed press-locking beyond autocorrelation in *JS* but not in *non-JS* trials, points the same way. A test of synchrony would need the partners’ signals in common trial time, which we did not analyse, and there the shorter delay in *JS* trials would itself change how much the partners’ windows overlap.

Respiratory depth was also aligned more strongly between partners in *JS* than in *non-JS* trials before the response (Fig 2B), at *t* = −12 and −6 s (uncorrected; the false discovery rate correction across the five windows gave *q* = 0.088 and 0.078), in the same anticipatory period in which the infra-slow EEG alignment was strongest (Fig 2A). Respiration modulates the brainstem, driving systemic shifts in attention and perception. Shared breathing depth before a joint response is therefore a candidate route through which the internal states of two partners converge, alongside the cardiac and electrodermal coupling described next.

The cardiac vagal index was aligned more strongly between partners in *JS* than in *non-JS* trials at *t* = − 6 s (Fig 2C), the window in which respiratory depth and the infra-slow EEG also aligned. This points to a parasympathetic route to a shared state in addition to the respiratory one, although it rests on a single window and needs replication. Skin conductance, a sympathetic measure, also aligned in an outcome-dependent way (Fig 2D, E). Accounts of social interaction give bodily signals a central role in alignment [52, 53], and these results suggest that social interaction engages peripheral sympathetic as well as parasympathetic dynamics [54, 52].

Our within-participant analyses (Fig 3) did not show reliable differences in brain–body coupling between successful and unsuccessful recognition. Coupling between the infra-slow EEG and the cardiac vagal index, between respiratory depth and the cardiac vagal index, and between the infra-slow EEG and tonic EDA (Fig 3D) did not differ between *JS* and *non-JS* at any window, and coupling with respiratory depth differed only at *t* = 0, uncorrected. The cardiac vagal index reduces the shape of a heartbeat scatter plot to a single nonlinear proxy for parasympathetic (vagal) activity [41], so these results give no evidence that vagal control, as indexed by this measure, was linked to the infra-slow EEG differently in successful and unsuccessful recognition. This model tests how a participant’s EEG tracks that participant’s own physiological signal within each window, so the result does not exclude coupling at other timescales. Previous research has demonstrated that coupling between infra-slow neural rhythms and autonomic processes occurs at different time scales [13, 55, 15]. Illustrative of this, a recent study on joint co-fluctuations of global fMRI activity with physiological signals used cross-correlation to demonstrate two distinct temporal relationships: a negative-to-zero lag peak (− 5 to 0 s), suggesting physiological signals follow global fMRI fluctuations, and a positive lag peak (0–10 s), indicating physiological signals may precede fMRI responses [29]. These bidirectional lags may point to the integration of top-down cortical modulation with bottom-up peripheral signalling, possibly reflecting system-wide autonomic regulation beyond local vascular effects [5, 29]. Overall, the inter-participant effects point to a role for infra-slow fluctuations in coordinating brain and bodily signals across partners, whereas the within-participant results give little evidence that this coupling depends on task outcome.

The one within-participant difference we observed, between respiratory depth and the motor-regressed infraslow EEG (0.08–0.12 Hz) at *t* = 0 (Fig 3A), is tentative: it is uncorrected for the five windows. It points, however, in the same direction as the inter-participant results. The body’s autonomic state, indexed here by breathing depth, and the infra-slow EEG rhythm may co-modulate each other, giving a state of readiness and sensory processing that anticipates the choice. On this reading, Fig 2A,B and Fig 3A may reflect the slow adjustment of gas exchange to the metabolic demands of attention, with tighter coupling between respiration and slow brain rhythms regulating the brain’s arousal state. High task performance occurs when this brain–body loop keeps the individual in an optimal attentional zone, and stronger coupling between respiratory depth and infra-slow EEG activity during optimal task execution reflects enhanced brain–body synchronisation that coordinates cortical excitability to support cognitive performance. Explanations of this kind remain highly speculative: our design cannot establish whether breathing drives the EEG, the EEG drives breathing, or a third process drives both, and a replication that specifies the *t* = 0 window in advance is needed to test it.

A supplementary analysis of phase at the click, pooled across all channels (Fig S2A, middle), showed a global, scalp-wide tendency of responses towards a phase of the *∼* 0.1 Hz wave. This tendency is descriptive, because channels at one click are not independent, and with one value per click phases did not deviate from uniformity in either condition (Supplementary Results). If it were found for individual responses, such a preference would suggest that infra-slow oscillations define temporal windows that optimise social coordination, similar to how faster neural oscillations create optimal windows for perception and action [46, 3]. A complementary within-participant analysis of the respiratory cycle at the click (Fig S2B) showed JS responses concentrated in the second half of expiration while non-JS responses concentrated in the first half of expiration. One possibility is that viscerosensory information modulates perception and cognition through temporal congruency, thereby colouring them with affective, value-related, and motivational significance [56, 5, 57, 58]. Interoceptive inference accounts make a related claim: bodily signals help shape the overall affective meaning of perception and action and general cognition [59]. According to this view, individuals with greater interoceptive abilities may perform better and show enhanced recognition in memory and social tasks [56]. In summary, we found that, at the dyadic level, participants who achieved mutual recognition (JS responses) showed significantly greater alignment in slow neuronal oscillations, the cardiac vagal index and phasic skin conductance compared with those who did not; alignment in respiratory depth and tonic skin conductance was also greater, but did not survive the correction across windows.

The respiratory and cardiac results can be read within a framework of cardiac–respiratory gating of joint decisions, in which two windows mark a quiet bodily state through different mechanisms. Expiration is a vagal-efferent window: respiratory sinus arrhythmia, as estimated by the cardiac vagal index, indicates that vagal outflow to the heart is greatest during expiration [60, 61]. Late diastole is a baroreceptor-quiescent window: arterial baroreceptors fire during systole and fall silent in diastole, so late diastole carries little cardio-vascular afferent signal, which does not by itself imply high parasympathetic tone [57, 62]. *JS* presses fell in the second half of expiration (Fig S2B), a pattern shared with voluntary actions in general, which are initiated more often during expiration [63]. This is not a general advantage of expiration, because memory and other cognitive functions have been tied to inspiration [64, 65]. Expiration may instead favour calm, internally paced acts such as a joint engagement, whereas inspiration promotes vigilant, encoding-heavy processing, in line with previous findings [66].

The cardiac phase at the press did not concentrate in late diastole, an expected finding, as self-initiated sampling has even been found to increase during systole [67], so a late-diastole window remains a potential hypothesis here. The cardiac result we did observe is positive but underpowered: at *JS* presses the phase of the infra-slow EEG was coupled with the cardiac phase, and this coupling differed from *non-JS* presses (Fig S3B; uncorrected). Neurovisceral integration models link vagally mediated cardiac control to flexible, regulated behaviour through shared cortical circuits [68], and in interoceptive inference accounts, low interoceptive prediction error could free precision for exteroceptive and social evidence [59, 69]. Neural responses to heartbeats, which predict perceptual decisions [70], may offer a cortical route to test this. Our results call for replication in larger samples.

### 4.2 Broader implications for social cognition and embodied interaction

Our findings have broader implications for understanding the neural basis of social cognition and embodied interaction [49]. Traditional approaches have focused on fast oscillations and event-related potentials, yet our results suggest that slower rhythms may also play an important role in organizing social interactions over extended timescales. Although the precise contribution of visceral and physiological components to perception and cognition still remains controversial, the alignment of infra-slow fluctuations between individuals (Fig 2A) may represent a previously overlooked mechanism for establishing shared cognitive and physiological states necessary for successful interpersonal coordination. This perspective builds on theories emphasizing the embodied and interactive nature of social cognition [22, 71, 51], and suggests that the very slow dynamics of neural activity provide a temporal scaffold upon which more rapid cognitive processes unfold.

A recent study using EEG and magnetoencephalography (MEG) to study the dynamics of intentional behavior has suggested that error detection arises from the interplay of two types of signals: a fast unconscious motor code, based on direct sensory and motor pathways, and a slower motor conscious intention that computes the required motor response given the stimulus and task instructions [47]. The supplementary tendency of correct task responses towards a phase of the *∼* 0.1 Hz wave (Fig S2A, middle), if confirmed, would extend this dual-route model of slow conscious versus fast nonconscious evidence accumulation to the infra-slow timescale, and suggest a role for slower visceral nonconscious components in shaping intentional behaviour. Their role would be to provide a slower nonconscious physiological context for the interplay of slow-conscious and fastnonconscious codes during the shaping of intentional and social behaviour.

Slow amplitude modulation of faster oscillations, which has been shown to be a mechanism for the integration of sensory information [46], could also help explain how infra-slow rhythms facilitate the temporal alignment implied during successful mutual coordination (Fig 2A). However, further research is needed to elucidate the precise mechanisms underlying this relationship and how it may vary across different social contexts and modalities. As noted, the brain’s amplitude modulation patterns crucial in shaping perception [72] and action may not be unique to the brain but extend to slower timescales of bodily physiological rhythms that entangle the internal and external milieus in service of adaptive and efficient social behavior. Our findings suggest that infra-slow rhythms may serve a role in social cognition by providing a temporal framework for coordinating complex interactions between individuals [3].

Future research should investigate whether these patterns generalize to other forms of social interaction and whether they can predict social outcomes in clinical populations with social impairments [73]. Methodological advances enabling the simultaneous measurement of infra-slow brain activity and physiological processes across interacting individuals will be crucial for further understanding the multilevel coordination that characterizes successful social interaction. By elucidating the role of infra-slow EEG rhythms in social cognition, we may prompt new ways of looking into the neural mechanisms underlying shared understanding and joint coordination, with implications for both basic and clinical research in social neuroscience.

### 4.3 Methodological considerations

A major challenge in studying infra-slow brain oscillations lies in conventional EEG methodology. Standard recordings typically implement DC-rejection or highpass filtering at the level of hardware, before digitisation, often attenuating or eliminating signals below 0.2 Hz [74]. To overcome this limitation, we employed DC-coupled amplifiers with ultra low cutoff (0.016 Hz) (see Methods). A second consideration is the imbalance of the data, which has two sources: there were more *JS* than *non-JS* trials, and dyads contributed unequal numbers of trials (Figure S1). We handled both within the APIM linear mixed-effects model and did not resample to equal counts, which would have discarded most of the *JS* trials. The model uses every eligible trial, so the smaller number of *non-JS* trials enters as greater uncertainty in that condition’s estimate, and its by-dyad random intercept and partner slope pool the estimates partially across dyads, so that dyads with many trials do not dominate the fixed effects (Supplementary Methods). The combined test across windows treats the five windows as independent although the signals are autocorrelated, so it should be read together with the window estimates (Table S1).

The perceptual crossing experiment offers important advantages for studying social alignment and embodied interaction. By reducing interaction to a minimalist sensorimotor loop, it isolates the temporal dynamics of interpersonal coordination while minimizing linguistic, visual, and cultural confounds [75]. This controlled simplicity enables the investigation of whether neuronal and physiological alignment emerges from a minimal interaction itself rather than from explicit communication or shared sensory input. Moreover, the continuous and reciprocal nature of the task allows the study of real-time mutual engagement and co-regulation of signals across multiple timescales. At the same time, the paradigm has limitations. Its abstract and constrained environment reduces ecological validity, and successful performance may partly reflect shared motor strategies or task structure rather than purely social processes. Thus, caution is warranted when generalizing these findings to more naturalistic social interactions.

Several limitations qualify our interpretation. The first principal component of infra-slow power explained most of the variance and had loadings of the same sign across the scalp, so it indexes a global fluctuation. Phase is less well defined for a component pooled across the scalp than for a single source, because it depends on the weights and the sign of the projection; for this reason, the coupling of this phase with the respiratory and cardiac cycles is reported in the Supplementary Results (Fig S3). We cannot exclude a contribution from global physiological or non-neural sources [76], such as skin potentials or ocular activity, although task trials were performed with eyes closed. Eye closure does not silence visual cortex, however, because haptic exploration in the dark can recruit retinotopic visual cortex [77]. The alignment effect did not depend on the frontopolar, anterior-frontal and rim channels most exposed to these sources (Supplementary Results). Cardiac activity also influences perception [78]. Coupling of this component with respiratory depth, tonic EDA and the cardiac vagal index was small (Fig 3), but its phase coupling with the cardiac cycle at the response differed with task outcome (Fig S3B), so a cardiac contribution cannot be excluded. The button press marks each participant’s decision to report recognition, which follows the first moment of recognition by an unknown delay. We use the term anticipatory for windows that precede the press, so alignment in those windows precedes the report of recognition and may not precede recognition itself. Finally, we did not relate infra-slow activity to participants’ ratings of perceptual awareness, so the link between individual perception and infra-slow activity remains to be tested.

### 4.4 Conclusion

Our findings suggest that the ∼0.1 Hz infra-slow EEG may provide a temporal scaffold for interpersonal coordination during real-time interaction. Mutual recognition was associated with enhanced inter-participant alignment in infra-slow EEG and autonomic signals. Infra-slow EEG and cardiac phases were coupled for successful but not unsuccessful recognition, a positive but underpowered result that calls for replication. Overall, interpersonal alignment at infra-slow timescales may offer a window onto the coordinated brain-body processes underlying shared understanding and mutual recognition.

## Acknowledgements

We warmly thank Tom Froese, who conceived the first drafts of this study and shaped its early direction. Professorial commitments prevented him from carrying the project to completion, but its conceptual foundation is his, and we are grateful for his generosity in entrusting these ideas to us. We also thank Nicolò Formento for writing code for the preprocessing.

## Author contributions

N. G. analyzed the data, adapted the theoretical and biological rationale for the APIM analysis across brain and physiological modalities following initial insights on the use and implications of the PCE paradigm provided by Tom Froese, and wrote the manuscript. S.G. extended the analysis to heart and respiration coupling and extended the interpretation and discussion of autonomic correlates of cognitive and physiological regulation. D.V. took the leading supervisory role, managed the funding and carried out the statistical checks. S.G. and D.V. revised the original draft, audited the code analyses and wrote the manuscript.

## Use of AI tools

The research questions, analysis rationale and analysis code were defined by the authors. Claude Code (Anthropic) was used to debug indexing issues in the original in-house scripts (Matlab and Python) and to check orthographic, syntactical and spelling issues during typesetting (format LATEX and .docx files). The authors reviewed and verified all AI-assisted code and text and take full responsibility for the content.

## Data and materials availability

The data used in this study is available from OSF at https://osf.io/84wbf/. The code used to analyze the data is available upon request from the corresponding author.

## Supplementary materials

### S1 Supplementary Methods

#### S1.1 Recording and preprocessing

Participants were asked to refrain from hair products the day before and from alcohol and caffeine in the 24 h preceding the session. Pair assignments were kept blind to the participants prior to the experiment (except for three pairs who were acquaintances or couples). Fpz was used as the ground electrode and FCz as the common reference. Electrode impedances were brought below 25 k*Ω* through conductive-gel preparation. A 60 Hz notch filter was applied to attenuate line noise in Okinawa. ECG was recorded with three electrodes (one below each clavicle and one over the lower-left rib cage) in a 3-lead configuration. Respiration was recorded with a piezo-electric belt placed on the upper abdomen. Electrodermal activity (EDA, skin conductance) was recorded via two electrodes placed on the tips of the index and ring fingers of the hand used to operate the button.

For the marking of transient artefacts, the MAD is preferred over the standard deviation for outlier detection in non-Gaussian, artefact-contaminated signals because it is not itself inflated by the outliers it is meant to flag [55], and the template of local-median rejection followed by spatial interpolation is a standard automated EEG artefact-rejection strategy [80, 81]. The 0.08–0.12 Hz band-pass also removes slow drifts, so no further detrending was applied. For the spatial whitening, we used the channel-level covariance data across its total length (rest, pause and task). This step scales the magnitude of the different sensor types relative to one another [82], effectively helping in the recovery of spatial patterns in noisy data with different magnitudes and artefacts across channels.

#### S1.2 Actor–Partner Interdependence Model

The model included partner, condition, and their interaction as fixed effects, plus a by-dyad random intercept and a random slope for partner, with dyad as the grouping variable. More explicitly, the model is fit, for each window, on a stacked observation set where each row corresponds to one participant-as-*actor* in one trial; because the APIM is symmetric, both members of each dyad contribute as *actor* once and as *partner* once per trial, so the design has *2 × N_dyad × K_trial* rows. The outcome vector y contains the actor’s window-averaged infra-slow EEG power; the design matrix X has four fixed-effect columns: an intercept (all ones), the partner’s window-averaged infra-slow

EEG power at the same window (continuous predictor), a binary condition indicator (0 = *non-JS*, 1 = *JS* ) attached to that trial, and their product (*partner × condition*). Regressing y on X yields four fixed-effect coefficients (*β*_0_, *β*_*partner, β*_*condition, β*_*partner×condition*); the reported *partner* × *condition* slope is therefore the difference, between *JS* and *non-JS*, in how strongly the partner’s infra-slow EEG power predicts the actor’s at that window — i.e., a per-window estimate of how mutual recognition modulates interpersonal alignment. The (*partner* | dyad_id) random structure adds a dyad-specific intercept and a dyad-specific partner slope, so that the fixed-effect estimates are pooled across dyads under partial pooling rather than dominated by individual pairs. The alignment plotted in the figures is *β*_partner_ for *non-JS* and *β*_partner_ + *β*_partner*×*condition_ for *JS*. Before fitting, the first principal component was scaled to the range 0–1 within each trial and within each dyad, and then z-scored within each participant over all trials and time points. Normalising over the whole epoch keeps the differences in power between windows and between conditions, which normalising within each analysis window would remove. We quantified alignment with regression because the slope measures how strongly one partner’s power predicts the other’s at each window. The model is symmetric: each participant enters once as actor and once as partner, so no assignment of participants to the x and y axes is required.

Three assumptions bear on the tests when the two conditions have unequal numbers of trials. The model has one residual variance for both conditions. The *p* value of each window is taken from a *t* distribution on the residual degrees of freedom of the model (the number of rows minus the four fixed effects; 408 for the EEG), without a small-sample correction, so it is close to a normal approximation. Stouffer’s Z combines the five windows as if they were independent. For the EEG, the onset-shift null provides a check that does not rest on these assumptions.

#### S1.3 Respiratory and cardiac signals

The respiratory cycle was derived from the raw respiratory belt signal by combining peak and trough detection [83] with a phase-completion estimator. The resulting respiratory phase (binary: 1 = inspiration, 0 = expiration) and respiratory cycle completion (0 → 1 within each half-cycle) were combined into a single full-cycle index in [0, 1].

This phase-completion representation captures the within-cycle position of each sample relative to the breathing rhythm and is well suited to coupling analyses with the ∼0.1 Hz infra-slow EEG band. Respiratory depth was computed from the cleaned belt signal: the squared signal was averaged over a centred 3 s running window, about one breath, and a shape-preserving cubic interpolant through the maxima of that average, at least 3 s apart, gave the upper envelope.

R peaks were detected in the cleaned ECG sampled at 100 Hz, and T offsets were taken from its delineation. For the cardiac phase, beats without a T offset, or with a systole shorter than 0.15 s or longer than 0.55 s (about 1% of beats), were excluded, and trials with a heart rate outside 30–200 bpm were excluded.

#### S1.4 Within-participant linear mixed-effects model

This within-participant approach is appropriate because (i) interoceptive and arousal mechanisms operate at the individual level, (ii) pooling participants (P1 and P2) increases statistical power while preserving trial structure, and (iii) there is no a priori reason to expect one partner’s EEG to predict the other’s autonomic state beyond shared environmental context, which is already captured in the dyadic analyses. The within-participant model predicted EEG from the physiological signal (respiratory depth, tonic EDA or the cardiac vagal index), condition, and their interaction, with a by-participant random intercept and a random slope for the physiological signal. The coupling plotted in the figures is *β*_physio_ for *non-JS* and *β*_physio_ + *β*_physio*×*condition_ for *JS*. The model of Figure 3C had the cardiac vagal index as the outcome and respiratory depth as the predictor, with the same specification and no EEG.

To compute the cardiac vagal index, we followed NeuroKit2 [40]: the index is log_10_(*L* × *T* ) [41], with *L* = 4 SD2 and *T* = 4 SD1, the longitudinal and transverse variability of the Poincaré plot of successive NN intervals. R peaks were detected in the cleaned ECG at 100 Hz. Trials with fewer than three R peaks or with a heart rate outside 30–200 bpm in either participant were excluded. A window used the NN intervals overlapping it. It was left out if it held fewer than three intervals, if any interval lay more than 20% from their median, or if SD1 or SD2 was zero. In the windowed model, the index computed over 5 s around each window centre was paired with the EEG power of the whole 6 s window.

#### S1.5 Phase at the click

To complement the dyadic analyses, we examined the relationship between the phase of the infra-slow EEG fluctuations (∼0.1 Hz) and task outcome at the individual level. Signal processing followed the procedure described in the Methods. EEG data were band-pass filtered in the 0.08 – 0.12 Hz range, and the Hilbert transform was applied to extract instantaneous phase at the timestamp of each button press. Phase values from all 64 channels were retained for subsequent circular statistics. Because instantaneous phase estimates based on the Hilbert transform are known to be inaccurate [84], we performed Monte Carlo simulations with synthetic signals of known phase. A sine wave at 0.1 Hz was contaminated with pink noise (1/f spectrum) at three signal-to-noise ratios relative to the amplitude of the sinusoid (noise amplitudes: 0.3, 0.5, 1.0) and processed using the same filtering and Hilbert transform steps that were applied to the empirical data. Phase accuracy was assessed at four reference phases (0°, 90°, 180°, 270°) across 90,000 measurements per condition (30 independent repeats). Results revealed a small systematic positive bias (9.8°–14.0°) consistent across conditions, with expected absolute errors of 11.0°–22.8°. The 0.08–0.12 Hz band yielded 95% CIs of ±13.3° to ±38.9°, providing interpretive context for systematic biases affecting empirical phase estimates.

The phases at the sample of each button press were pooled across all 64 channels and all participants and grouped by task outcome (19,328 values from the 302 *JS* clicks and 7,040 from the 110 *non-JS* clicks). Circular histograms (four bins of 90°, one for each quarter of the cycle) quantified phase distributions, expressed as the change from the mean bin share in percentage points. Rayleigh tests of deviation from circular uniformity were applied separately to each condition. Because the channels at one click are not independent, these tests describe a global, scalp-wide tendency, and their *p* values should be read descriptively. For the bars of Figure S2 we also tested each bin for more responses than chance, in that direction only, with an exact one-sided binomial test (the channel-pooled EEG counts, the EEG with one value per click, and the respiratory cycle, with chance of one quarter per bin). The four segments of the cardiac cycle differ in duration, because systole fills about 40% of a beat but a quarter of the phase axis, so for each click chance was the share of the samples of its own trial and participant that fell in each segment, and the count of a bin was tested exactly as the sum of these click-wise probabilities (Poisson-binomial). The four one-sided *p* values of each panel and condition were corrected with the Benjamini–Hochberg procedure (false discovery rate 0.05).

#### S1.6 Bivariate von Mises sine model

For two phases at the click, written here for respiratory phase *ϕ*_*r*_ and infra-slow EEG phase *ϕ*_*b*_, the model is

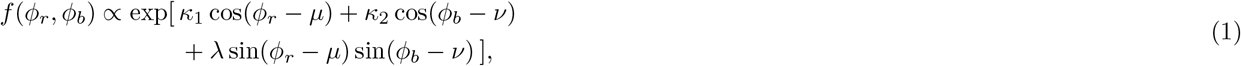

where *µ* and *v* are the mean phases, *κ*_1_ and *κ*_2_ the concentrations, and *λ* the coupling between the phases. We fitted the model separately to *JS* and *non-JS* clicks by maximum likelihood, restricted to unimodal solutions. EEG phase *ϕ*_*b*_ was the phase of the motor-regressed 0.08–0.12 Hz analytic signal at the click, projected onto the first principal component of the power around the click and sign-aligned on the motor channels. For the pairs that include the EEG, we built the null distribution with the same surrogate procedure, re-reading both phases at each surrogate sample.

To locate differences in the phase plane, we evaluated the fitted densities on a 60 × 60 grid. Clusters were connected grid cells, with circular wrap-around, whose value exceeded the 85th percentile of the pooled absolute null surface. For the density of each condition, clusters were computed on the observed density minus the mean surrogate density. Cluster mass was the sum of the absolute values over the cluster, and cluster *p* values came from the null distribution of the maximum cluster mass. No cluster was significant (smallest *p* = 0.144). As a test that does not depend on the fitted model, we also counted the clicks of each condition in the 16 combinations of two quarter-cycle bins (4 × 4 test). The bins were the quarters starting at 0, 90, 180 and 270 degrees for the respiratory phase (0 degrees is the trough, 180 the peak) and the EEG phase (0 degrees is the peak, 180 the trough), and the four segments (systole and early, mid and late diastole) for the cardiac phase. Chance was taken from the first 50 surrogate sets, with both phases re-extracted at the surrogate click times, so that the unequal time spent in each bin was accounted for. For each cell, we tested whether the share of clicks exceeded that of the surrogates and, for the difference, whether the share was larger for *JS* than for *non-JS* clicks (one-sided); the *p* value was the proportion of surrogate sets with a value at least as large as observed. To correct across the 16 cells, the z score of each cell (observed minus surrogate mean, divided by the surrogate standard deviation) was compared with the largest z score among the 16 cells of each surrogate set. With 50 sets, the smallest attainable *p* value is 0.020.

### S2 Supplementary Results

#### S2.1 Spectral peak at 0.1 Hz

To test whether the task EEG contained power around 0.1 Hz above the 1/f background, we computed a multitaper spectrum (3 Slepian tapers) [35] of each 60-second trial and channel, and averaged it over channels and over each participant’s trials. We fitted a straight line to log power against log frequency, leaving out 0.07–0.13 Hz. The residual from this line was averaged in the 0.08–0.12 Hz band and tested against zero across the 62 participants. Power at 0.08–0.12 Hz exceeded the background (mean residual +0.200 log_10_ units, SD 0.147; *t*(61) = 10.70, *p* < 0.001; sign-flip permutation *p* < 0.001), and 94% of participants were above the fit. In a control band of 0.14–0.18 Hz, the residual did not differ from zero (mean −0.003, *t*(61) = −0.55).

#### S2.2 Trial-count balance across dyads

**Fig. S1.**
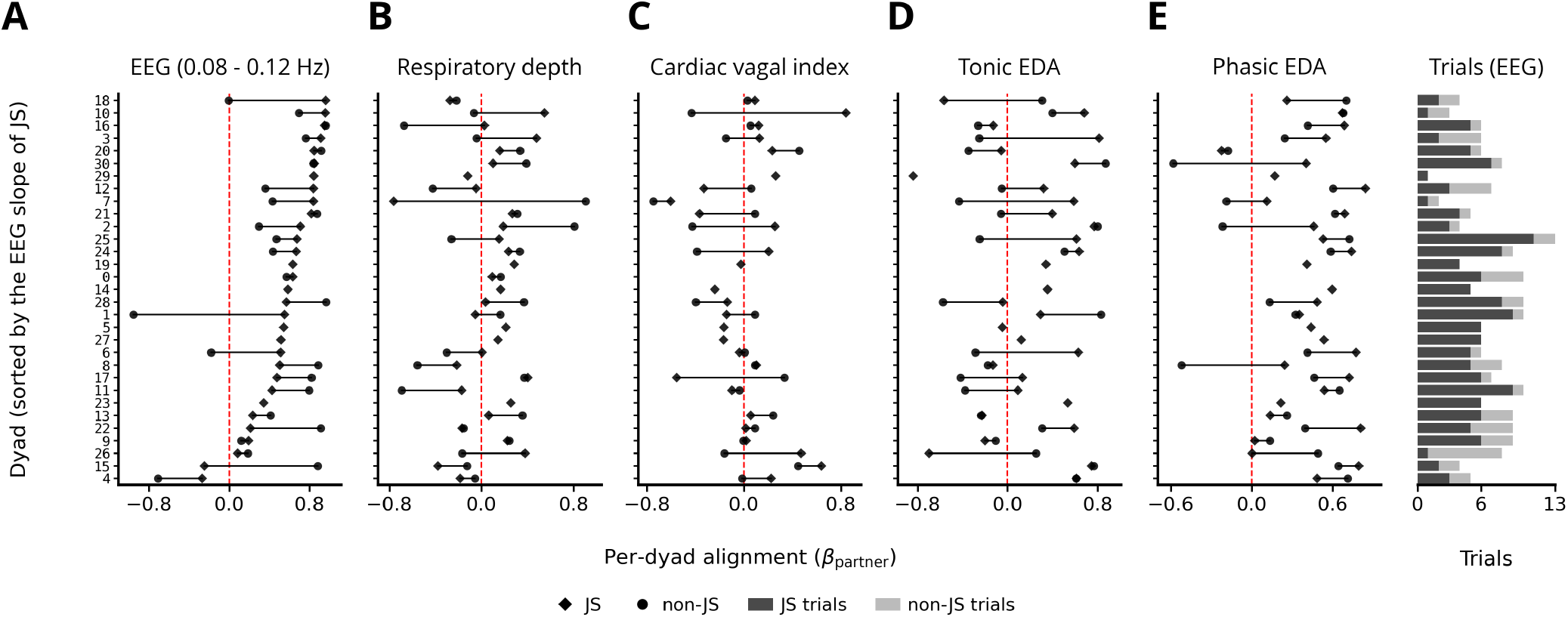
Dyad-level ordinary least-squares assessment of trial-count balance. Per-dyad alignment for *JS* (diamonds) and *non-JS* (circles), joined by a line for each dyad. Per-dyad alignment is the slope of actor on partner from an ordinary least-squares fit without intercept, pooled across windows (*β*_partner_). Every panel has the dyads in the same order, sorted by the *JS* slope of the EEG, so that a dyad keeps its row; a dyad without data for a signal leaves an empty row. Red dashed line, zero. **(A)** The 0.08–0.12 Hz EEG band (31 dyads). **(B)** Respiratory depth (31 dyads). **(C)** Cardiac vagal index (27 dyads). **(D)** Tonic and **(E)** phasic skin conductance (30 dyads). Right: trials per dyad in the EEG analysis, stacked by outcome; C to E use subsets of these trials.

To check that the task-level effects did not rest on a few dyads or on unequal trial counts, we computed for each dyad and condition the slope of actor on partner from an ordinary least-squares fit without intercept, pooled across the five windows (Figure S1). The EEG and respiratory depth analyses included 151 *JS* and 55 *non-JS* trials from 31 dyads, the cardiac vagal index 121 and 48 trials from 27 dyads, and the electrodermal signals 145 and 51 trials from 30 dyads. In the EEG analysis, dyads contributed between 1 and 13 trials (median 6), and 25 of the 31 dyads had trials of both outcomes; the other six had *JS* trials only. These per-dyad slopes are descriptive: inference rests on the mixed-effects models, which pool the dyads partially (Supplementary Methods).

#### S2.3 Inter-participant alignment with a channel subset

To test whether the inter-participant effect in Figure 2A depended on the channels most exposed to ocular and other non-neural sources, we computed the first principal component without 15 frontopolar, anterior-frontal and rim channels (FP1, FP2, AF7, AF3, AFz, AF4, AF8, F7, F8, FT9, FT10, T7, T8, TP9 and TP10) and refitted the same model; this analysis is reported here without a figure. With the remaining 49 channels, alignment was larger in *JS* than *non-JS* across windows (Stouffer Z = 2.44, *p* = 0.007; one-sided *p* = 0.039, 0.015, 0.327, 0.434 and 0.180 at *t* = −12, −6, 0, +6 and +12 s). With all 64 channels the result was Z = 2.00, *p* = 0.023, and with the 15 excluded channels alone it was Z = 1.40, *p* = 0.081. The trends were largely the same as with all channels (Figure 2A), with one extra significant window: at *t* = −12 s the difference reached significance with 49 channels (*p* = 0.039) but not with 64 (*p* = 0.350). The window at −6 s remained significant (*p* = 0.015, against 0.003 with 64 channels), and the windows at 0, +6 and +12 s stayed non-significant in both analyses.

#### S2.4 Electrodermal activity

We extended the dyadic APIM to the tonic skin conductance level (Fig 2D) and the phasic skin conductance response (Fig 2E), at the same five windows (N=145 *JS* and N=51 *non-JS* trials for both). Tonic EDA alignment was significantly stronger in *JS* than *non-JS* at *t* = −12 s and *t* = +6 s (one-sided, uncorrected; neither survived the false discovery rate correction across the five windows, *q* = 0.17), which may reflect a slow build-up and recovery of shared autonomic tone around the task event. Phasic EDA alignment separated the groups in the complementary windows at *t* = −6 s (*JS β* = 0.10, 95% CI −0.05 to 0.25; *non-JS β* = −0.17, 95% CI −0.38 to 0.03; difference 0.28, 95% CI 0.05 to 0.50) and *t* = +12 s (*JS β* = 0.24, 95% CI 0.11 to 0.37; *non-JS β* = −0.10, 95% CI 0.34 to 0.13; difference 0.35, 95% CI 0.08 to 0.62; both survived the false discovery rate correction, *q* = 0.043), suggesting that the phasic component tracks more transient co-fluctuations flanking the response. The two electrodermal components therefore contributed to inter-participant alignment on distinct, interleaved timescales rather than a single locked-in window. Within participants (Fig 3D), coupling of tonic EDA with motor-regressed EEG did not differ between *JS* and *non-JS* at any window (smallest one-sided *p* = 0.130). Without the motor regression the smallest one-sided *p* was lower but still not significant (0.070, at *t* = −12 s, non-*JS* > *JS* ; not shown), so removing the pre-movement motor potential reduced the evidence for a difference in coupling.

#### S2.5 Phase at the click

To assess the channel-pooled phase preference in Figure S2A, middle without treating channels as independent observations, we reduced the phases to one value per click. With the circular mean over all 64 channels (Figure S2A, right), phases did not deviate from uniformity in *JS* (*z* = 0.55, *p* = 0.58, mean phase 174°, mean resultant length 0.04) or in *non-JS* (*z* = 1.26, *p* = 0.28, mean phase 64°, mean resultant length 0.11). The channel-pooled tendency is therefore not supported at the level of individual clicks. Without the motor regression, the channel-pooled distributions were similar (Figure S2A, left; *JS* : *z* = 8.4, *p* < 0.001, mean phase 188°, mean resultant length 0.02; *non-JS* : *z* = 20.3, *p* < 0.001, mean phase 76°, mean resultant length 0.05; both peaked in the 90–180° bin), so the pooled tendency does not depend on the motor regression.

**Fig. S2.**
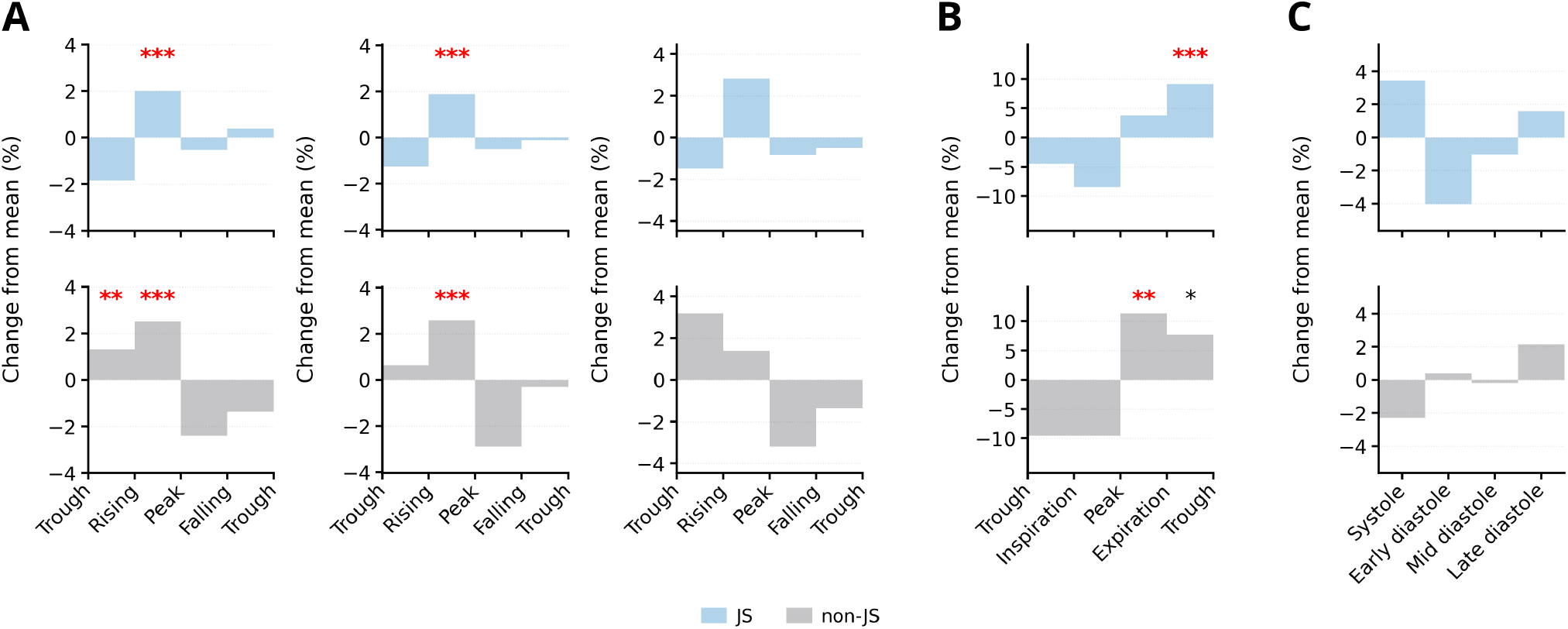
Phase of the infra-slow EEG, respiration and the heart at the response. Change from the mean, in percentage points, in four phase bins at the click, for *JS* (upper row of each panel, blue; N=302 clicks, N=270 in C) and *non-JS* (lower row, grey; N=110 clicks, N=103 in C). Asterisks mark bins above chance (one-sided test, uncorrected): * *p* < 0.05, ** *p* < 0.01, *** *p* < 0.001; bold red asterisks also survive the correction across the four bins of the panel (false discovery rate 0.05). **(A)** The 0.08–0.12 Hz EEG band. Left and middle, the phases of all 64 channels pooled at each click (19,328 *JS* and 7,040 *non-JS* values), from the EEG that was not motor-regressed (left) and from the motor-regressed EEG (middle); the pooled values show a global, scalp-wide tendency, and because the channels at one click are not independent, these tests are descriptive. Right, one value per click, the circular mean over all 64 channels of the motor-regressed EEG. **(B)** The respiratory cycle (Trough → Inspiration → Peak → Expiration). **(C)** The cardiac cycle (Systole → Early, Mid and Late diastole), where the mean is the share expected from the time spent in each segment.

#### S2.6 Phase coupling of the infra-slow EEG with the respiratory and cardiac cycles

We asked whether the coupling between the phase of the respiratory cycle and of the infra-slow EEG at the moment of the response differed with task outcome. Figure S3A shows the fitted density of the two phases for *JS* and *non-JS* clicks and their difference. Phase coupling did not differ significantly with task outcome: *λ* was +0.259 for *JS* (*p* = 0.085) and −0.272 for *non-JS* (*p* = 0.418), and the difference did not reach significance (*Δλ* = +0.531, *p* = 0.085, onset-shift null). In the 4 × 4 test, clicks gathered in the cell that combines late expiration (the last quarter of the respiratory cycle) with the late rising phase of the EEG wave (the quarter before its peak), in both conditions (*JS* : 10.3% of clicks against 7.5% in the surrogates, *p* = 0.039; *non-JS* : 11.8% against 6.8%, *p* = 0.020). Neither cell survived the correction across the 16 cells (corrected *p* = 0.451 and 0.314), and no cell held a larger share of *JS* than of *non-JS* clicks (smallest *p* = 0.098).

We next asked whether the cardiac cycle was coupled with the phase of the infra-slow EEG at the click (Figure S3B). At *JS* presses, the phase of the infra-slow EEG was coupled with the cardiac phase (*λ* = +0.303, *p* = 0.030), whereas at *non-JS* presses it was not (*λ* = −0.366, *p* = 0.184), and the coupling differed between conditions (*Δλ* = +0.669, *p* = 0.040, onset-shift null, uncorrected; Figure S3B). In the 4 × 4 test, no cell held more clicks than chance in either condition, or a larger share of *JS* than of *non-JS* clicks (smallest uncorrected *p* = 0.098), so the difference concerned the overall coupling of the two phases rather than a specific combination of them.

**Fig. S3.**
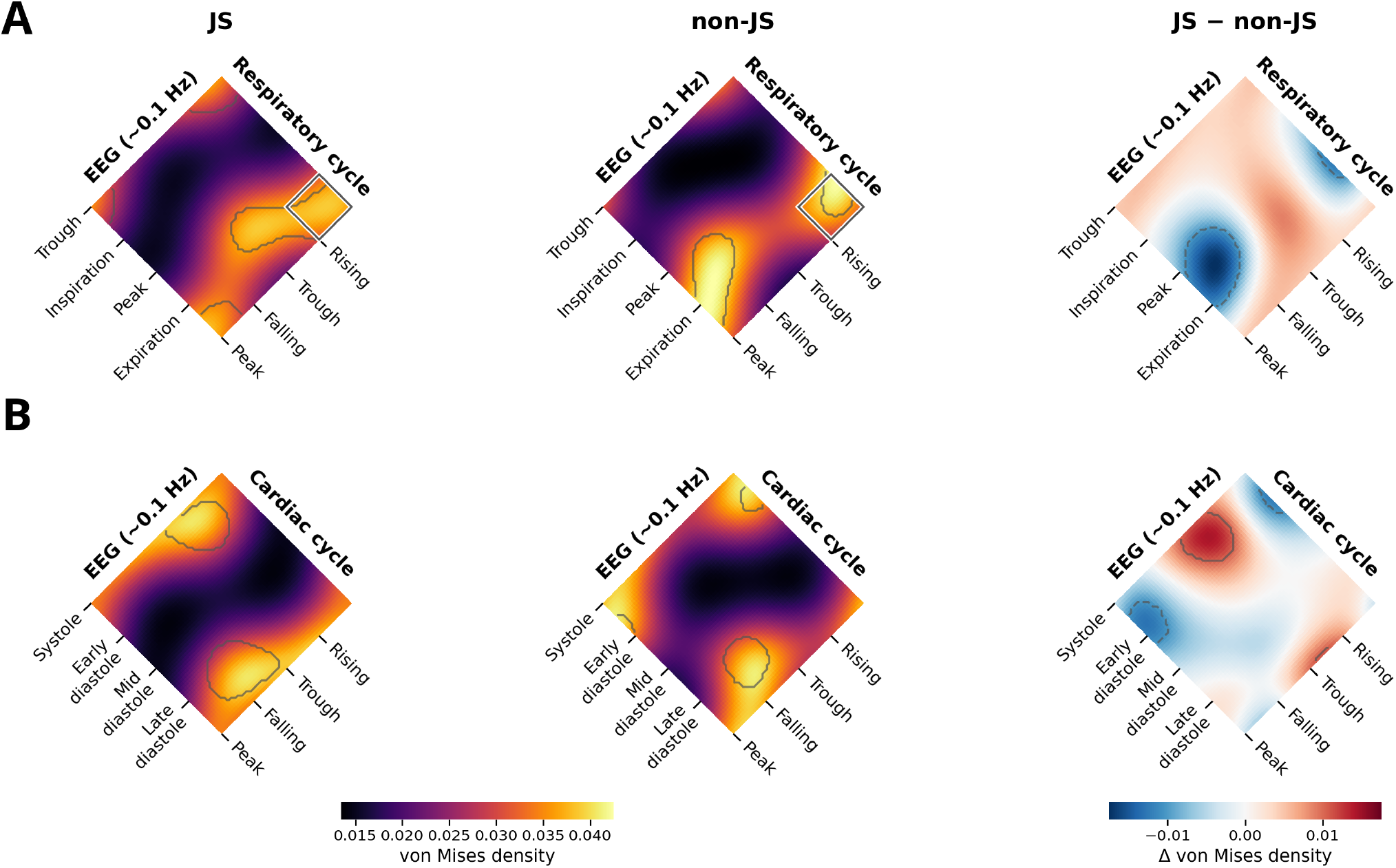
Phase coupling of the infra-slow EEG with the respiratory and cardiac cycles at the click. Fitted bivariate von Mises densities for *JS, non-JS* and their difference, as labelled above each column. Both rows share one colour scale for the densities and one for the difference. Curved outlines mark clusters above the 85th percentile of the onset-shift null: in the *JS* and *non-JS* columns, clusters of density above the mean of the null; in the difference column, clusters of the difference (solid, positive; dashed, negative). Black outlines, *p* < 0.05 in the cluster test; grey outlines, not significant (all). Square outlines mark cells of the 4 × 4 test, in which clicks were counted in the 16 combinations of two quarter-cycle bins and compared with the onset-shift null (one-sided): in the *JS* and *non-JS* columns, cells with more clicks than chance; in the difference column, cells with a larger share of *JS* than of *non-JS* clicks. Grey squares, *p* < 0.05 uncorrected; black squares, significant after correction across the 16 cells (none). Labels and tick marks give the position of each landmark on its axis: the trough and peak of the respiratory cycle with mid-inspiration and mid-expiration between them, the peak and trough of the EEG wave with the midpoints of its falling and rising flanks, and the centre of each cardiac segment. Cardiac phase runs through systole and early, mid and late diastole (Methods); because systole fills a larger share of each beat than of the axis, density gathers in systole even when clicks are spread evenly in time. **(A)** Respiratory cycle phase and motor-regressed infra-slow EEG phase (N=302 *JS* and N=110 *non-JS* clicks). **(B)** Motor-regressed infra-slow EEG phase and cardiac phase (N=270 *JS* and N=103 *non-JS* clicks).

#### S2.7 Window estimates of the mixed-effects models

Table S1 gives, for every panel of Figures 2 and 3 and every window, the slope of each condition and the difference between conditions, with their 95% confidence intervals, and the uncorrected and corrected *p* values. The signals were normalised before fitting, so the slopes are in standard-deviation units and can be read as effect sizes.

**Table S1:** Window estimates of the mixed-effects models of Figures 2 and 3. *β*, slope of each condition with its 95% confidence interval; *Δβ*, difference between *JS* and *non-JS* (the interaction term of the model) with its standard error (SE) and 95% confidence interval (estimate ± 1.96 SE). *p*, uncorrected: one-sided for *JS* > *non-JS* in Figure 2A–C; the smaller of the two one-sided values in Figure 2D, E and Figure 3, where both directions were tested. *q*, after the Benjamini–Hochberg correction across the five windows of a panel (from twice the smaller one-sided *p* where both directions were tested). Bold, *p* or *q* < 0.05; heavier bold, *q* < 0.05 (significant after correction).

| Window (s) | *JS β* [95% CI] | *non-JS β* [95% CI] | *Δβ* (SE) [95% CI] | *p* | *q* |
|---|---|---|---|---|---|
| *Figure 2A: EEG* |  |  |  |  |  |
| −12 | 0.03 [−0.11, 0.17] | −0.01 [−0.23, 0.21] | 0.04 (0.11) [−0.18, 0.27] | 0.350 | 0.449 |
| −6 | 0.17 [0.02, 0.32] | −0.13 [−0.34, 0.07] | 0.31 (0.11) [0.09, 0.53] | **0.003** | **0.016** |
| 0 | 0.13 [0.00, 0.27] | −0.02 [−0.22, 0.19] | 0.15 (0.12) [−0.08, 0.38] | 0.102 | 0.254 |
| +6 | 0.10 [−0.09, 0.30] | 0.14 [−0.13, 0.41] | −0.03 (0.12) [−0.28, 0.21] | 0.605 | 0.605 |
| +12 | 0.02 [−0.14, 0.18] | −0.03 [−0.29, 0.24] | 0.05 (0.13) [−0.21, 0.31] | 0.359 | 0.449 |
| *Figure 2B: respiratory depth* |  |  |  |  |  |
| −12 | 0.07 [−0.07, 0.21] | −0.14 [−0.35, 0.08] | 0.21 (0.12) [−0.02, 0.44] | **0.035** | 0.088 |
| −6 | 0.08 [−0.03, 0.20] | −0.16 [−0.36, 0.03] | 0.25 (0.12) [0.02, 0.48] | **0.016** | 0.078 |
| 0 | 0.00 [−0.10, 0.11] | −0.01 [−0.20, 0.18] | 0.01 (0.11) [−0.21, 0.23] | 0.465 | 0.557 |
| +6 | −0.11 [−0.23, 0.01] | −0.18 [−0.39, 0.02] | 0.08 (0.12) [−0.16, 0.31] | 0.263 | 0.438 |
| +12 | −0.08 [−0.22, 0.06] | −0.07 [−0.28, 0.15] | −0.02 (0.12) [−0.25, 0.22] | 0.557 | 0.557 |
| *Figure 2C: cardiac vagal index* |  |  |  |  |  |
| −12 | −0.04 [−0.19, 0.12] | 0.05 [−0.19, 0.29] | −0.09 (0.13) [−0.35, 0.17] | 0.744 | 0.852 |
| −6 | 0.16 [0.03, 0.28] | −0.35 [−0.53, −0.17] | 0.51 (0.12) [0.27, 0.74] | **< 0.001** | **< 0.001** |
| 0 | −0.24 [−0.37, −0.12] | −0.16 [−0.43, 0.10] | −0.08 (0.15) [−0.38, 0.21] | 0.710 | 0.852 |
| +6 | −0.07 [−0.20, 0.06] | −0.16 [−0.36, 0.05] | 0.09 (0.13) [−0.16, 0.34] | 0.249 | 0.621 |
| +12 | −0.17 [−0.36, 0.02] | −0.02 [−0.29, 0.25] | −0.15 (0.14) [−0.42, 0.13] | 0.852 | 0.852 |
| *Figure 2D: tonic EDA* |  |  |  |  |  |
| −12 | 0.19 [0.02, 0.36] | −0.01 [−0.22, 0.20] | 0.20 (0.11) [−0.01, 0.42] | **0.034** | 0.170 |
| −6 | 0.21 [0.06, 0.37] | 0.07 [−0.12, 0.26] | 0.14 (0.11) [−0.07, 0.36] | 0.098 | 0.326 |
| 0 | 0.10 [−0.05, 0.25] | 0.07 [−0.14, 0.29] | 0.03 (0.12) [−0.20, 0.26] | 0.404 | 0.905 |
| +6 | 0.01 [−0.16, 0.18] | −0.21 [−0.43, 0.02] | 0.21 (0.12) [−0.01, 0.44] | **0.033** | 0.170 |
| +12 | −0.11 [−0.26, 0.04] | −0.09 [−0.32, 0.13] | −0.01 (0.12) [−0.24, 0.21] | 0.453 | 0.905 |
| *Figure 2E: phasic EDA* |  |  |  |  |  |
| −12 | −0.04 [−0.21, 0.12] | −0.05 [−0.23, 0.13] | 0.01 (0.12) [−0.22, 0.23] | 0.473 | 0.946 |
| −6 | 0.10 [−0.05, 0.25] | −0.17 [−0.38, 0.03] | 0.28 (0.12) [0.05, 0.50] | **0.009** | **0.043** |
| 0 | −0.09 [−0.23, 0.06] | 0.04 [−0.17, 0.24] | −0.12 (0.11) [−0.35, 0.10] | 0.137 | 0.457 |
| +6 | −0.01 [−0.19, 0.17] | 0.01 [−0.21, 0.23] | −0.02 (0.11) [−0.24, 0.21] | 0.445 | 0.946 |
| +12 | 0.24 [0.11, 0.37] | −0.10 [−0.34, 0.13] | 0.35 (0.14) [0.08, 0.62] | **0.006** | **0.043** |
| *Figure 3A: respiratory depth and EEG* |  |  |  |  |  |
| −12 | 0.00 [−0.14, 0.13] | 0.02 [−0.21, 0.25] | −0.03 (0.13) [−0.29, 0.24] | 0.424 | 0.898 |
| −6 | −0.04 [−0.15, 0.08] | −0.01 [−0.19, 0.17] | −0.03 (0.10) [−0.23, 0.18] | 0.402 | 0.898 |
| 0 | 0.13 [0.01, 0.25] | −0.05 [−0.24, 0.13] | 0.18 (0.11) [−0.03, 0.39] | **0.047** | 0.475 |
| +6 | 0.15 [−0.01, 0.32] | 0.00 [−0.24, 0.24] | 0.15 (0.13) [−0.11, 0.42] | 0.123 | 0.614 |
| +12 | 0.08 [−0.04, 0.21] | 0.10 [−0.10, 0.30] | −0.02 (0.12) [−0.25, 0.22] | 0.449 | 0.898 |
| *Figure 3B: cardiac vagal index and EEG* |  |  |  |  |  |
| −12 | −0.03 [−0.15, 0.09] | −0.01 [−0.21, 0.18] | −0.02 (0.12) [−0.24, 0.21] | 0.442 | 0.988 |
| −6 | −0.04 [−0.14, 0.07] | −0.04 [−0.18, 0.11] | 0.00 (0.09) [−0.18, 0.17] | 0.494 | 0.988 |
| 0 | 0.00 [−0.11, 0.10] | −0.10 [−0.33, 0.13] | 0.10 (0.13) [−0.15, 0.35] | 0.219 | 0.988 |
| +6 | −0.03 [−0.21, 0.16] | −0.01 [−0.28, 0.25] | −0.01 (0.14) [−0.29, 0.27] | 0.464 | 0.988 |
| +12 | −0.07 [−0.20, 0.06] | 0.05 [−0.17, 0.27] | −0.12 (0.13) [−0.37, 0.13] | 0.180 | 0.988 |
| *Figure 3C: respiratory depth and cardiac vagal index* |  |  |  |  |  |
| −12 | 0.10 [−0.09, 0.30] | 0.26 [−0.03, 0.55] | −0.16 (0.17) [−0.48, 0.17] | 0.172 | 0.887 |
| −6 | 0.10 [−0.09, 0.29] | 0.01 [−0.27, 0.30] | 0.09 (0.16) [−0.23, 0.40] | 0.295 | 0.969 |
| 0 | −0.02 [−0.18, 0.15] | 0.01 [−0.23, 0.24] | −0.03 (0.14) [−0.30, 0.25] | 0.427 | 0.969 |
| +6 | 0.06 [−0.13, 0.24] | 0.06 [−0.20, 0.32] | −0.01 (0.15) [−0.30, 0.29] | 0.485 | 0.969 |
| +12 | 0.09 [−0.06, 0.24] | 0.22 [0.00, 0.43] | −0.13 (0.14) [−0.39, 0.14] | 0.177 | 0.887 |
| *Figure 3D: tonic EDA and EEG* |  |  |  |  |  |
| −12 | −0.03 [−0.16, 0.10] | 0.02 [−0.15, 0.19] | −0.04 (0.10) [−0.24, 0.15] | 0.328 | 0.696 |
| −6 | −0.01 [−0.10, 0.09] | −0.04 [−0.18, 0.10] | 0.03 (0.08) [−0.13, 0.20] | 0.348 | 0.696 |
| 0 | −0.03 [−0.18, 0.11] | −0.09 [−0.31, 0.13] | 0.06 (0.14) [−0.22, 0.33] | 0.340 | 0.696 |
| +6 | −0.03 [−0.15, 0.09] | −0.09 [−0.30, 0.12] | 0.06 (0.12) [−0.18, 0.30] | 0.313 | 0.696 |
| +12 | −0.02 [−0.15, 0.12] | −0.17 [−0.40, 0.06] | 0.15 (0.13) [−0.11, 0.40] | 0.130 | 0.696 |

